# Low-dimensional brain-symptom associations delineate depression phenotypes with distinct connectivity biomarkers and symptom profiles

**DOI:** 10.1101/2025.08.19.670030

**Authors:** Wenya Liu, Maria Vesterinen, Alexandra Andersson, Paula Partanen, Samanta Knapič, Joonas J. Juvonen, Felix Siebenhühner, Antti Salonen, Hanna Renvall, Risto J. Ilmoniemi, Eero Castrén, Erkki Isometsä, Dimitri Van De Ville, J. Matias Palva, Satu Palva

## Abstract

Depression is neurobiologically and clinically heterogeneous. New approaches using resting-state functional MRI (rs-fMRI) functional connectivity (FC) data have modeled the neural basis of depression heterogeneity and revealed unique neural phenotypes. Yet, no studies have identified depression phenotypes from electrophysiological magnetoencephalography (MEG) data although MEG measures human brain dynamics at millisecond precision. We demonstrate here unique depression phenotypes based on MEG-oscillation FC. We collected resting-state MEG, MRI, and clinical symptom data from 263 patients with unipolar depression and 75 healthy controls. We assessed MEG-FC with two oscillatory coupling-mode measures that are fundamental for information processing. To define normative phenotypes, we computed their latent-space low-dimensional brain-symptom associations, and used these components to identify phenotypes using unsupervised machine learning. We identified five stable depression phenotypes that were characterized by unique symptom profiles and distinct spectral patterns. Our results demonstrate new neural underpinnings of depression heterogeneity and reveal unique neural phenotypes with potential personalized diagnostic value.

## Introduction

Major depressive disorder (MDD) is a highly heterogeneous disorder, encompassing a diverse range of cognitive and behavioral symptoms as well as complex comorbidities with other brain disorders (Goldberg, 2011; McQuaid, 2021). This clinical heterogeneity may contribute to the wide range of suicide risk factors (Holma et al., 2010). Depression heterogeneity complicates diagnosis and limits the efficacy and predictability of current treatments (Hyman, 2007). During the past decade, neuroimaging research has led to MDD being increasingly recognized as a system-level disorder of brain network activity, with abnormalities, *e.g.*, in functional connectivity (FC) (Chai et al., 2023; Gong & He, 2015; Kaiser et al., 2015; Spellman & Liston, 2020). Studies using resting-state functional magnetic resonance imaging (rs-fMRI) have consistently linked MDD to changes in the organization and dynamics of FC in the limbic and frontostriatal networks (Drysdale et al., 2017; Lynch et al., 2024; Touron et al., 2022), the default mode network (DMN) (Furman et al., 2011; Liston et al., 2014; Sheline, Barch, Price, Rundle, Vaishnavi, Snyder, et al., 2009; Zhou et al., 2020), or to the crosstalk between the networks (Kaiser et al., 2015; Mulders et al., 2015). Especially time-varying dynamic coupling appears to mediate distinct clinical symptoms (Piguet et al., 2021). While altered frontostriatal FC is linked to mood regulation, sustained negative affect, and rumination (Furman et al., 2011), the hypoconnectivity of the frontoparietal control network is associated with deficits in cognitive control (Schultz et al., 2019). Furthermore, increased FC between DMN and subgenual prefrontal cortex can reliably predict levels of depressive rumination (Hamilton et al., 2015; Siddiqi & Fox, 2024).

Mapping brain-symptom associations with multivariate approaches (Kebets et al., 2021; Vieira et al., 2024; Xia et al., 2018) and clustering subjects based on their co-occurring brain activity patterns (Griffa et al., 2022; Haakana et al., 2024; Van De Ville et al., 2021) is a clinically useful approach to understand the impact of brain dynamics heterogeneity on symptom presentation in mental disorders. Pioneering studies have combined symptom-specific fMRI-FC network data with machine learning (ML) methods and revealed biological depression phenotypes corresponding to distinct symptom profiles and responsivity to brain stimulation interventions (Chen et al., 2023; Drysdale et al., 2017; Dunlop et al., 2024; Liang et al., 2020; Price et al., 2017). More recently, normative models (Sun et al., 2023; Tozzi et al., 2024) and personalized brain scores (Sun et al., 2023; Tozzi et al., 2024) have identified depression phenotypes with different sensitivity to pharmacological interventions. Such findings have fueled the growing conceptualization of depression treatments as network therapies, aimed at modulating specific neural circuits to alleviate specific symptoms (Cash et al., 2021; Chai et al., 2023; Dunlop et al., 2019; Fox, 2018). On the other hand, a recent meta-analysis of normative findings did not find consistent generalization of individual-level biomarkers for MDD (Winter et al., 2024), thus necessitating new methods that could overcome the limitations in the current approaches.

While the fMRI signal reflects slow neuronal-activity related fluctuations in blood oxygenation in multi-second time scales (< 1 Hz), electro- and magnetoencephalography (EEG/MEG) directly measure neuronal population activity with millisecond precision (Baillet, 2017). This activity is characterized by rhythmic oscillations in sub-second temporal scales (> 1 Hz), which arise from the interactions between excitatory pyramidal neurons and GABAergic inhibitory interneurons. Oscillations are fundamental for neuronal computations and provide millisecond-resolution temporal clocking mechanisms by creating temporally-correlated windows of excitability (Engel et al., 2001; Fries, 2015; Singer, 1999; Thut et al., 2012). Transiently coupled large-scale oscillatory networks have been found to be essential for dynamic routing of information in healthy brain functions (Engel et al., 2013; S. Palva & Palva, 2012) and, conversely, aberrant oscillation dynamics is characteristic to neuropsychiatric disorders (Uhlhaas et al., 2017). Crucially, one key cellular-level pathophysiological factor in MDD is disrupted GABAergic inhibitory functioning and excitation/inhibition (E/I) imbalance associated with deficient somatostatin (SST)-interneuron-driven inhibition (Anderson et al., 2020; Fee et al., 2017). In line with this, local oscillatory activity as measured by EEG is aberrant at several frequency bands in depression (see reviews by (Fernández-Palleiro et al., 2020; Fingelkurts & Fingelkurts, 2015; Smart et al., 2015)), including interhemispheric asymmetry in the alpha band (Allen et al., 2004; Bruder et al., 1997), altered theta activity in anterior cingulate cortex (Pizzagalli, 2011; Pizzagalli et al., 2003), and increased beta power (Grin-Yatsenko et al., 2010). Altered oscillatory activity also shows plastic modulation to pharmacological and nonpharmacological treatments (Bruder et al., 2008; Korb et al., 2009; Smart et al., 2015). Source-reconstructed MEG data, with superior spatial accuracy over EEG, have further confirmed oscillatory alterations in depression, including abnormal gamma oscillations in the visual cortex (Dai et al., 2024) and irregular beta bursts in the orbitofrontal cortex (Xue et al., 2025). Recent EEG studies demonstrated the biomarker potential of local alpha and theta oscillations (W. Wu et al., 2020), alpha peak frequency (Li et al., 2025; Voetterl et al., 2023), and their relevance for intervention outcome prediction (Schwartzmann et al., 2024; Watts et al., 2022).

Despite the promising fMRI-FC findings, evidence of electrophysiological FC alterations in depression has remained scarce. A recent study found greater beta-2 band (18.5–21 Hz) connectivity within DMN, and greater beta-1 band (12.5–18 Hz) connectivity between the DMN and the frontoparietal network (FPN) (Miljevic et al., 2023; Saletu et al., 2010; Whitton et al., 2018), while another report found decreased alpha-band connectivity in depression (Nugent et al., 2020). Despite the promise of revealing dynamic coupling directly linkable to underlying neural mechanisms and neurophysiological inhibitory deficits via M/EEG recordings, no studies so far have leveraged oscillatory-coupling-based FC data to identify putative depression phenotypes by clustering patients according to their co-occurring oscillatory and synchronization patterns.

To address this knowledge gap, we collected symptom data using 10 self-report questionnaires (**Fig. 1 a**), resting-state MEG and magnetic resonance imaging (MRI) (**Fig. 1 b**) data from 263 patients with unipolar depression and from 75 healthy controls and computed individual brain oscillation connectomes (**Fig. 1 c–d**). We then computed multivariate brain-symptom associations (**Fig. 1 f**) and leveraged the latent components to identify depression phenotypes with clustering algorithms (**Fig. 1 g**).

**Figure 1.**
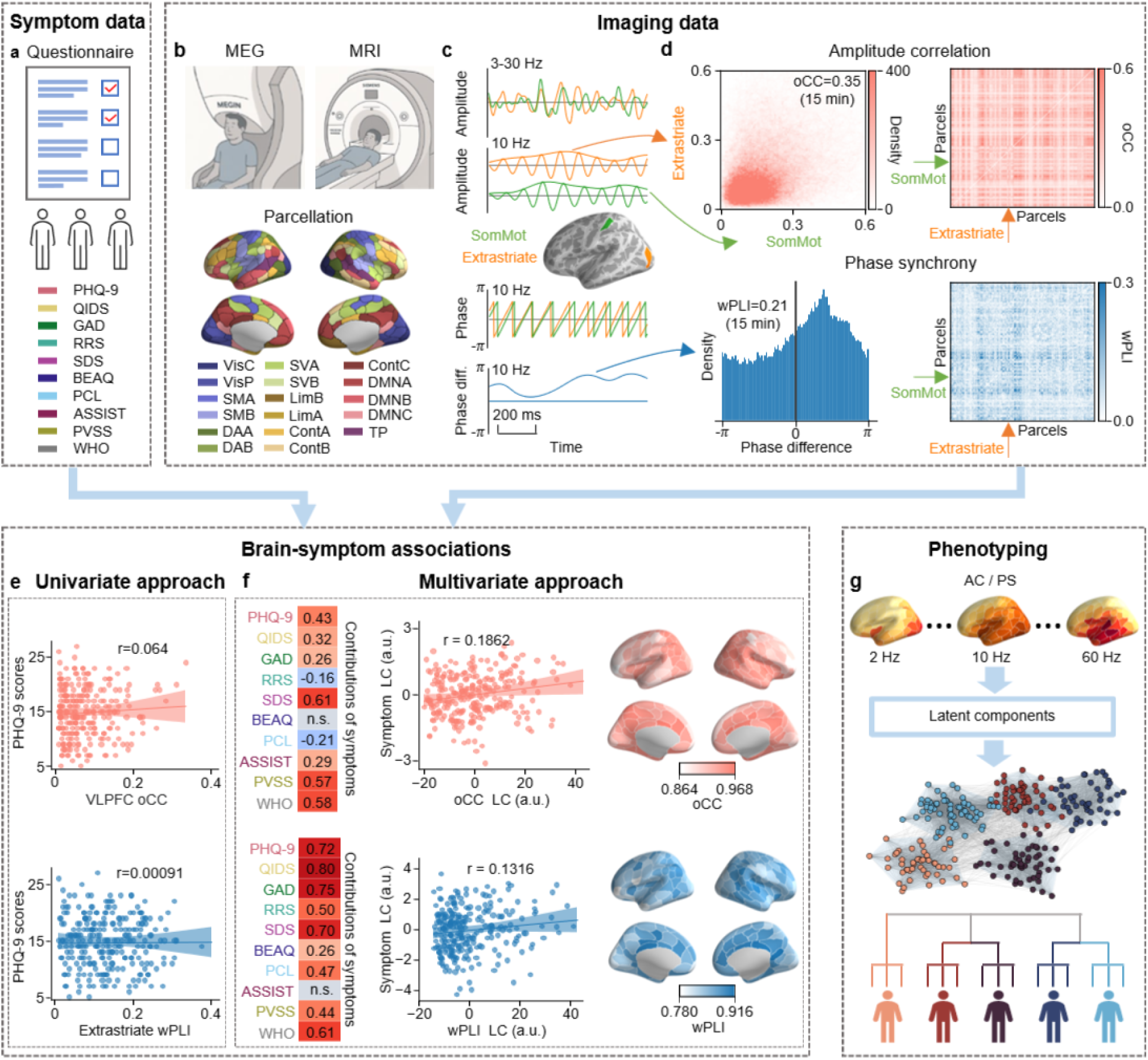
Schematic overview of the study. **a** Symptom data were collected with 10 self-report symptom scales: Patient Health Questionnaire (PHQ-9), Quick Inventory of Depressive Symptomatology 16-item (self-report) (QIDS), Generalized Anxiety Disorder 7-item (GAD), The Ruminative Responses Scale Short-version (RRS), Sheehan Disability Scale for functional impairment (SDS), Brief Experiential Avoidance Questionnaire (BEAQ), PTSD Checklist for DSM-5 (PCL), Alcohol, Smoking, and Substance Involvement Screening Test Lite (ASSIST), Positive Valence Systems Scale (PVSS), and WHO-5 Well-Being Index (WHO). **b** Magnetoencephalography (MEG) data were source-reconstructed using individual structural magnetic resonance imaging (MRI) and parcellated into the Schaefer atlas with 200 parcels encompassing 17 networks. Schematic pictures created with ChatGPT. **c** Broadband MEG time series (top row) for each parcel were filtered with a Morlet wavelet to obtain narrow-band amplitudes (second row), phase (third row), and phase difference (last row) time series. **d** Pairwise amplitude correlation (AC) was computed with orthogonalized correlation coefficient (oCC), and pairwise phase synchrony (PS) was assessed with weighted phase lag index (wPLI) from phase differences. **e** Brain-symptom associations were assessed with Pearson correlation, a univariate approach, for node centrality in both AC and PS, and for each symptom scale. **f** Brain-symptom associations were assessed with partial least squares correlation (PLSC), a multivariate approach, applied separately for AC and PS. The illustrated examples show PLSC results obtained with all symptom scales and AC/PS around 10 Hz, including the contribution of symptom scales (left column), Pearson correlations of latent components (middle column) from symptoms and node centrality of AC (top row) and PS (bottom row), and contribution of node centrality (right column). **g** PLSC was conducted on symptom data and AC/PS across 32 frequencies. Significant latent components of AC/PS node centrality were then combined as input for Leiden clustering to derive data-driven depression phenotypes.

## Results

### Symptom heterogeneity

The clinical cohort comprised 263 patients diagnosed with MDD. The presence of a depressive episode was verified with the Mini-International Neuropsychiatric Interview (M.I.N.I.) and patients were required to have an existing contact to the healthcare. Among the MDD cohort, with the mean age being 34.3 years (18–60 years), 62% (*N*=163) of patients were identified as females, 33% (*N*=87) as males, and 5% (*N*=13) as “other” (**Supplementary Figure S1 a**). No significant differences in age were found between any pair of gender categories (two-tailed Mann-Whitney U test, *p*=0.061 for male-female, *p*=0.536 for female-other, and *p*=0.155 for male-other comparisons). The healthy control (HC) cohort, with the mean age being 31.9 years, comprised *N*=75 participants with 39% (*N*=29) as females, 60% (*N*=45) as males, and 1% (*N*=1) as “other” (**Supplementary Figure S1 b**). No significant differences in age were found between male and female cohorts (two-tailed Mann-Whitney U test, *p*=0.458). The MDD and HC groups differed significantly in gender distribution (Chi-square test, χ^2^ =14.94, *p*=0.0001 for male-female comparison) and age (two-tailed Mann-Whitney U test, *p*=0.005).

We used the Patient Health Questionnaire (PHQ-9) as the primary measure of depression severity. The PHQ-9 scores for the clinical cohort (*N*=263) ranged from 5 to 27, with a mean of 14.82 and a standard deviation (SD) of 4.66. The clinical cohort comprised 15%, 31%, 36%, and 18% of patients with mild, moderate, moderately severe, and severe depression (thresholds: 5, 10, 15, and 20), respectively (**Fig. 2 a**). In addition to PHQ-9, in order to map the heterogeneity of depression, symptoms were collected with 9 other questionnaires: Quick Inventory of Depressive Symptomatology 16-item (self-report) (QIDS), Generalized Anxiety Disorder 7-item (GAD), The Ruminative Responses Scale Short-version (RRS), Sheehan Disability Scale (SDS), Brief Experiential Avoidance Questionnaire (BEAQ), PTSD Checklist for DSM-5 (PCL), Alcohol, Smoking, and Substance Involvement Screening Test Lite (ASSIST), Positive Valence Systems Scale (PVSS), and WHO-5 Well-Being Index (WHO). (see details in **Methods**). All 10 questionnaire scores were normalized to a 0–1 scale based on the minimum and maximum values of their native score range, and WHO and PVSS scores were inverted for better comparability with the other scores, so that a higher normalized score consistently indicated more severe symptoms (**Fig. 2**). All symptom scores had high heterogeneity across the depression cohort, the scores being the smallest for ASSIST measuring substance abuse, and the largest for the inverted WHO-5 scale indicating overall poor wellbeing in the cohort (**Fig. 2 a**). Females in the MDD cohort had significantly higher scores for depressive severity (QIDS), anxiety (GAD), rumination (RRS), and PTSD (PCL), and lower scores for anhedonia (PVSS) and substance abuse (ASSIST) than males (1,000 bootstrapping at the alpha level of 0.05, ***Supplementary Figure S2)***.

**Figure 2.**
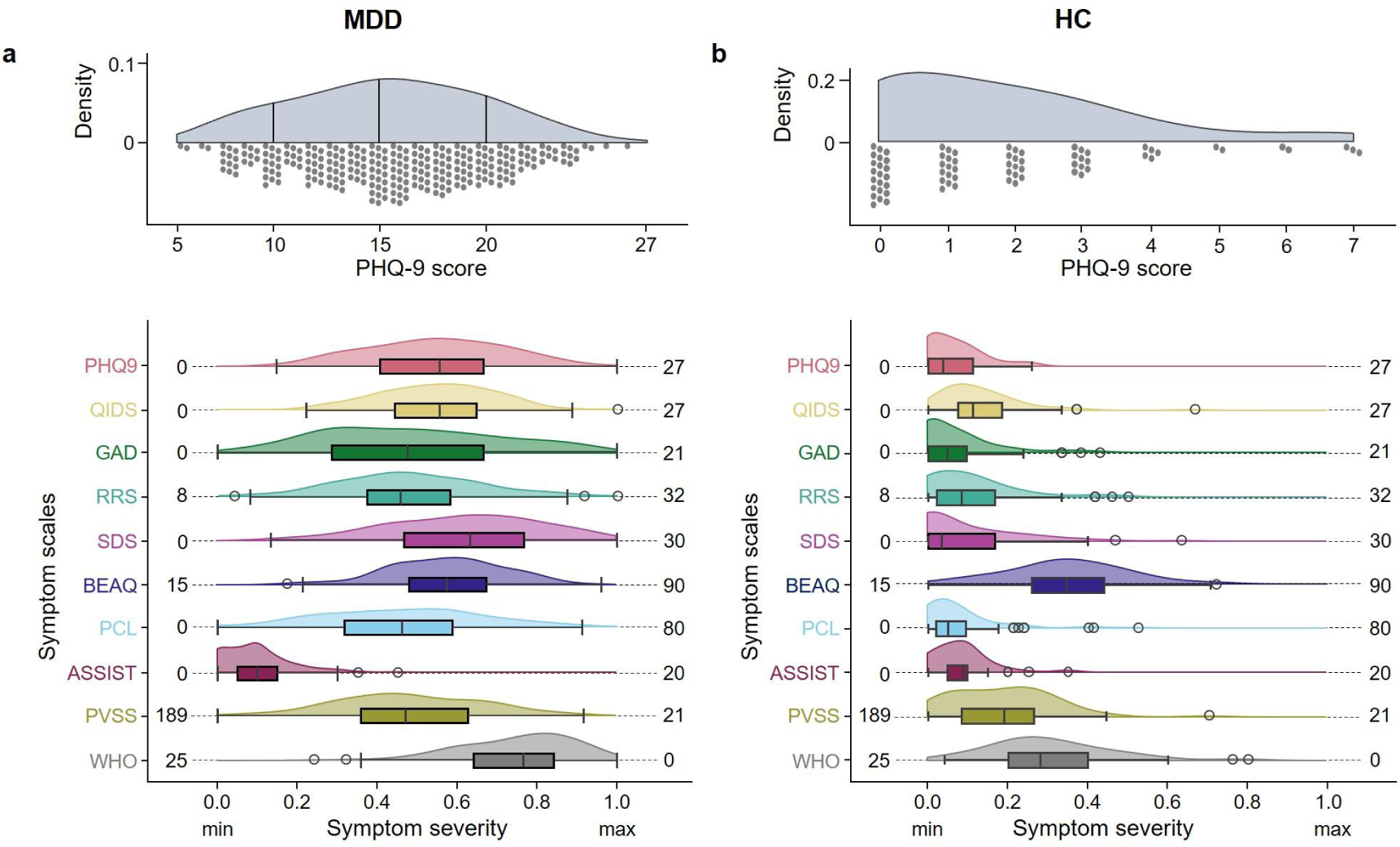
Symptom score distributions for the Major Depressive Disorder (MDD) cohort (a) and the healthy control (HC) cohort (b). Top panel: Distribution of PHQ-9 scores which assess the overall severity of depression. Total scores of 5, 10, 15, and 20 represent cutting points for mild, moderate, moderately severe and severe depression, respectively. Bottom panel: Distribution of all 10 symptom scores, using both boxplots and probability density functions (PDFs), including PHQ-9, QIDS, GAD, RRS, SDS, WHO, PVSS, BEAQ, PCL, and ASSIST. All scores were normalized to a 0–1 scale based on the minimum (left side numbers) and maximum (right side numbers) values of the native score range for each individual questionnaire. Please note that WHO and PVSS scores were inverted (subtracted from 1 after min-max normalization) to maintain the direction of symptom severity, ensuring that a higher normalized score consistently indicates more severe symptoms.

For the MDD cohort, 32.7% of patients had only been diagnosed with depression, while the rest had one or several comorbid diagnoses, most commonly anxiety (21.3%) or ADHD (6.8%) (**Supplementary Figure S1 c**). In the HC cohort, the symptom scores were significantly smaller compared to the MDD cohort and below the clinical cut-offs (**Fig. 2 b**). While 5 subjects had higher scores on PHQ-9, they were undiagnosed and didn’t fulfill the criteria of depression (M.I.N.I. module) before filling in self-report questionnaires, and they were therefore grouped to the HC group.

### Phase synchrony and amplitude correlation networks

The 15-min resting-state MEG data were source-reconstructed using individual MRI-derived head models, collapsed into 200 cortical parcels of the Schaefer atlas, and Morlet-filtered from 2-60 Hz. Oscillation amplitudes averaged across parcels peaked in the alpha band at 10 Hz as shown before (Haegens et al., 2014; Javed et al., 2025) and showed no significant differences between MDD and HC groups (**Supplementary Figure S3,** two-tailed Mann-Whitney U test, p<0.05, False Discovery Rate (FDR) corrected).

Oscillation-based connectivity was estimated at each frequency between all parcel pairs for two coupling modes, inter-areal phase synchrony (PS) and amplitude correlations (AC), which reflect partially dissociated and complementary mechanisms of neuronal interactions (Engel et al., 2013; Hipp et al., 2012; Siebenhühner et al., 2024; Siems & Siegel, 2020; Zhigalov et al., 2017). Inter-areal PS was estimated with the weighted phase lag index (wPLI) and AC with the orthogonalized correlation coefficient (oCC), which are insensitive to direct effects of source leakage (J. M. Palva et al., 2018). The oscillatory networks were represented as graphs with parcels as nodes and connections between parcels as edges (Bullmore & Sporns, 2009). At the whole-brain level, we estimated graph strength (GS) (the average of strength across all edges), which peaked in the alpha (8–13 Hz) band for both coupling modes as previously reported for resting-state data (Fuscà et al., 2023; Siebenhühner et al., 2024); the peak was wider for AC (5–30 Hz) than for PS (6–15 Hz) (**Fig. 3 a‒b**). No statistically significant differences were found between the HC and MDD cohorts in GS for either coupling mode (**Fig. 3 a‒b**, **Supplementary Figure S4,** two-tailed Mann-Whitney U test, *p*>0.05, FDR corrected). To map PS and AC connectivity at the parcel level, we computed node strength (NS) for each parcel. The MDD cohort exhibited weaker AC node strength in the delta band (2–4 Hz) compared to the HC cohort. In contrast, PS node strength was weaker in the theta (4‒5 Hz) and alpha-low-beta (10–15 Hz) bands in MDD than HC subjects, but stronger in the delta (2–4 Hz) and beta (19–21 Hz) bands (**Fig. 3 c‒d, Supplementary Figure S5**) underscoring only minor differences at the group-level as found before in neuroimaging (Winter et al., 2022). To understand if AC and PS convey similar information, we computed their correlations on graph strength. In both HC and MDD groups, PS and AC graph strength was correlated in the theta-alpha and beta bands, as previously observed in healthy subjects (<u>Siebenhühner et al., 2024</u>), and additionally in the gamma band in the MDD group (***Supplementary Figure S6***). Although significant, these correlations were only moderate indicating that the coupling modes yield complementary information.

**Figure 3.**
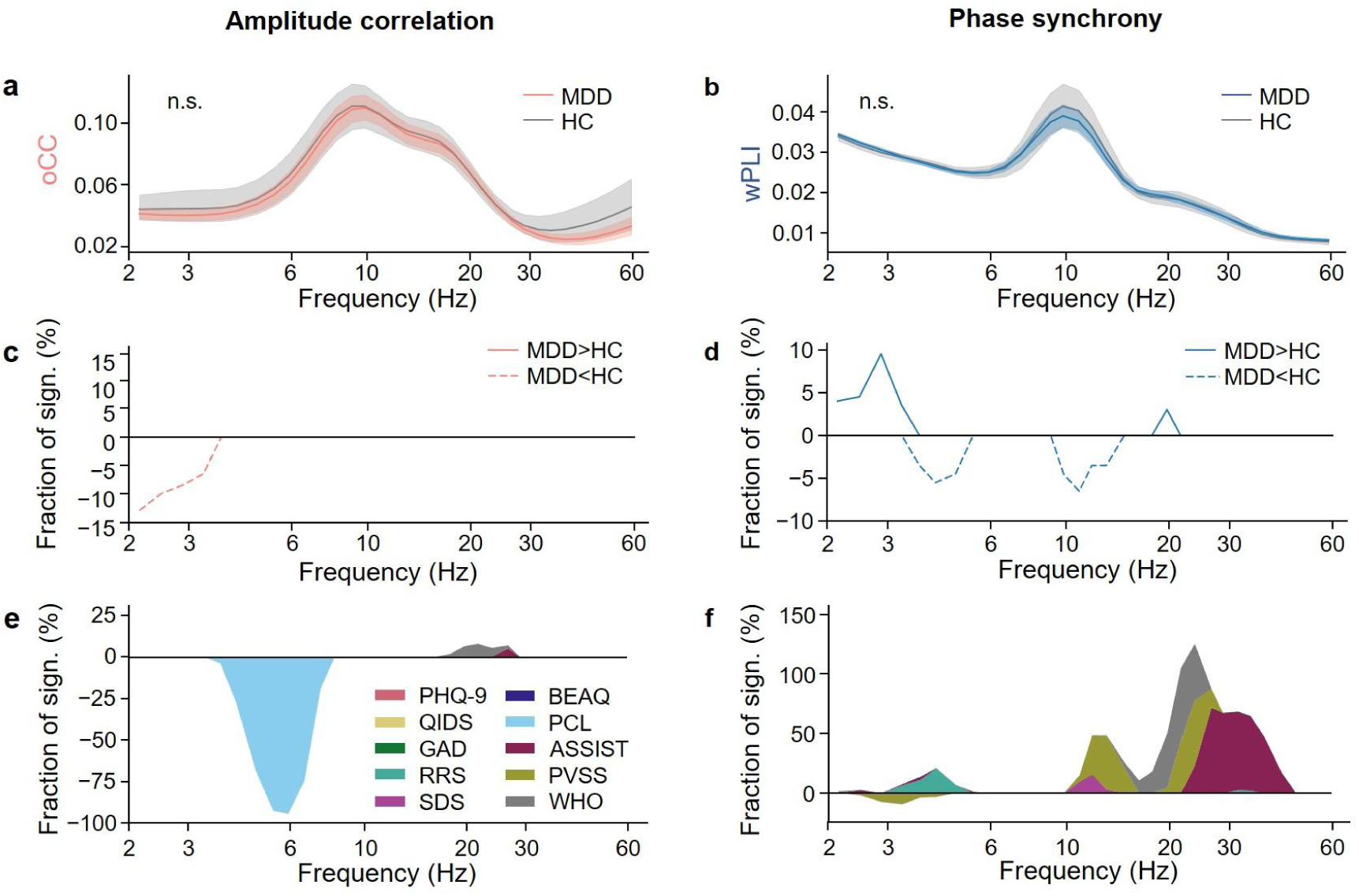
Amplitude correlations (AC), phase synchrony (PS), and their univariate associations with symptom scores. **a**–**b** The spectrum of graph strength of oCC (**a**) and wPLI (**b**) from the MDD cohort (N=263) and the healthy control (HC) cohort (N=75). No significant differences in graph strength were found for both AC and PS between MDD and HC cohorts (two-tailed Mann-Whitney U test, p<0.05, False Discovery Rate (FDR) corrected). **c–d** Fraction of parcels where MDD and HC cohorts differed significantly (two-tailed Mann-Whitney U test, alpha-level subtraction, 0.025 for each tail) for AC (**c**) and PS (**d**). **e–f** Stacked plots of the fraction of parcels that were significantly correlated with each symptom score (permutation-based multiple comparison correction at p<0.05, see details in Methods) for AC (**e**) and PS (**f**).

We next computed their association with symptom severity with univariate Pearson correlations between node strength and each symptom score (**Fig. 3e–f**). For AC, node strength in the theta band (4–7 Hz) showed significant correlations with the PCL scale assessing post-traumatic stress disorder (PTSD) symptoms. For PS, alpha band (here 12-15 Hz) node strength was correlated with PVSS measuring anhedonia, and beta band (15-28 Hz, and 22-50 Hz) with WHO assessing well-being and ASSIST measuring substance abuse, respectively. The multiple correlations across PS and AC suggested that rather than univariate approaches, multivariate approaches should be used to estimate brain-symptom correlations.

### Brain-symptom associations of phase synchrony and amplitude correlation networks

To further understand latent brain-symptom associations, we applied a multivariate approach using partial least squares correlation (PLSC). Age and gender effects were first regressed out, and we then calculated the cross-covariance of symptoms and node strength for AC and PS separately. To estimate the statistical significance of latent components, the singular values of the cross-covariance from the observed data were compared against 1,000 permuted values. One significant latent component was derived from AC and another from PS (**Fig. 4 a**). Correlations between the significant brain latent component and the symptom latent component were significant for both AC (**Fig. 4 b** top panel, *r*=0.1798, *p*=0.0034) and PS (**Fig. 4 b** bottom panel, *r*=0.2046, *p*=0.0008).

**Figure 4.**
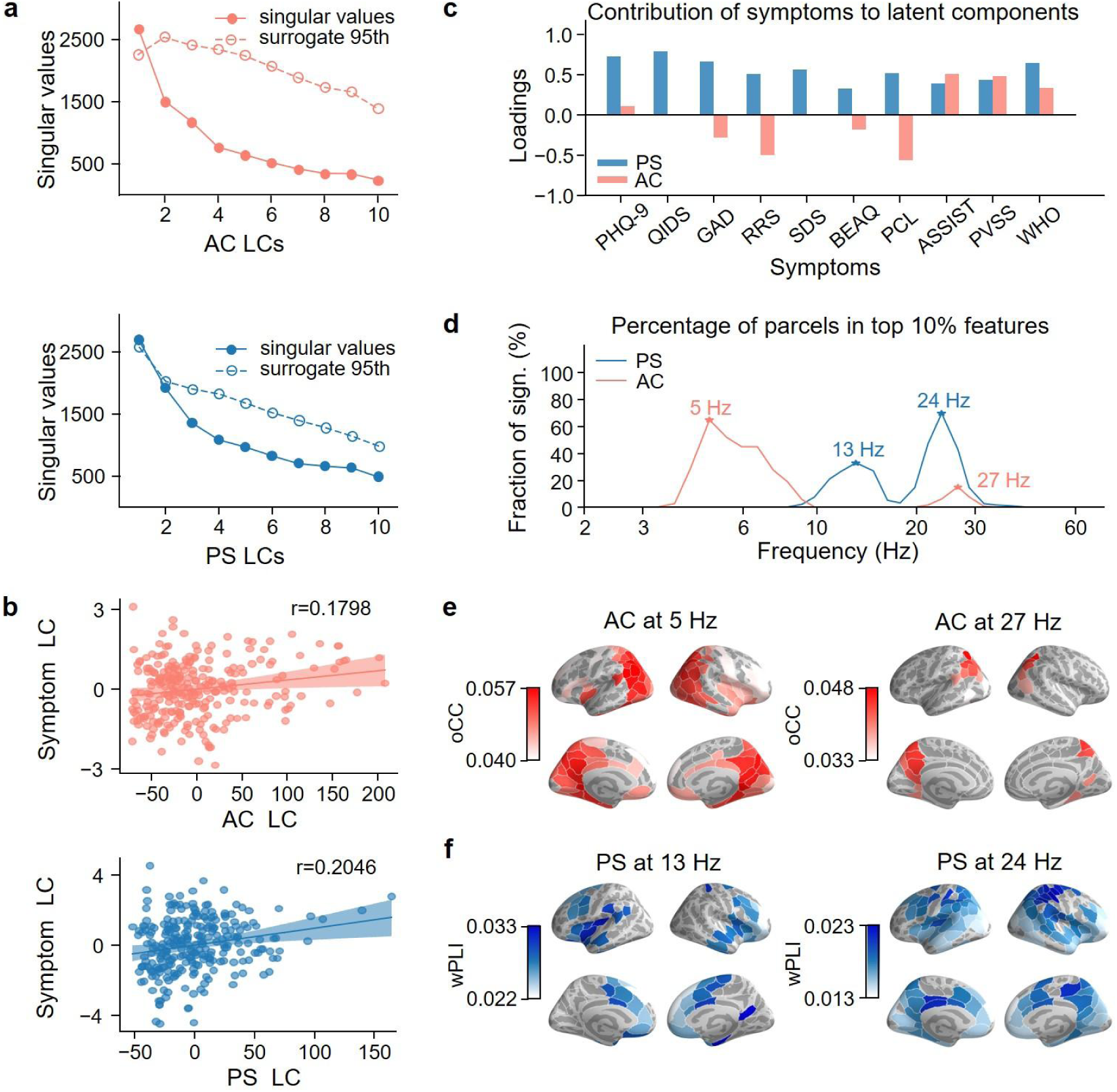
Brain-symptom associations with partial least squares correlation (PLSC) in MDD cohort. **a** Singular values from the singular value decomposition (SVD) of the cross-covariance matrix between oCC node centrality and symptom scores (top), and between wPLI node centrality and symptom scores (bottom). LC = Latent component. **b** The Pearson correlation between the symptom latent component and the node centrality latent component (top: amplitude correlation, bottom: phase synchrony). **c** The contribution of symptoms to the two significant latent components (one from PS, p=0.0452 and one from AC, p=0.0085). **d** Fraction of parcels per frequency among the top 10% of features contributing the most to the two latent components. **e** Spatial patterns of wPLI at two peak frequencies (13 and 24 Hz) in (**d**). **f** Spatial patterns of oCC at two peak frequencies (5 and 27 Hz) in (**d**).

The AC-derived latent component was associated with symptom dimensions including rumination (RRS), anhedonia (PVSS), post-traumatic stress disorder (PCL), substance abuse (ASSIST), and well-being (WHO) (**Fig. 4 c**). Symptom contributions to this latent component were distributed across both positive and negative loadings, underscoring differential and complementary symptom relevance with AC. In contrast, the PS-derived latent component captured the overall severity of depressive symptoms as measured with PHQ-9 and QIDS, along with the other 8 symptom scales (**Fig. 4 c**). The contributions of symptoms to the latent components were generalizable and stable, except for QIDS and SDS in the AC-derived components, whose contributions were minimal (A generalizability and stability test; ***Supplementary Figure S7 a***).

To reveal the spectral contributions to the latent components, we plotted the fraction of parcels per frequency among the top 10% node strength values which contributed the most to the latent components. For the AC-derived latent component, the key contributions emerged in theta (4–8 Hz) and beta (25–28 Hz) bands (**Fig. 4 d**). These spectral profiles were robust against the cut-off percentage (***Supplementary Figure S8***). The theta-band contributions were dominated by the ventral attention network (VAN) and the visual network, while the beta-band contributions were primarily influenced by the VAN, DMN, and the visual network (**Fig. 4 e**, ***Supplementary Figure S9 a, c***). In contrast, for the PS-derived latent component, the key contributions emerged in the alpha band (11–16 Hz) predominantly associated with VAN, and the beta band (22–26 Hz) primarily linked to DMN and the control network (**Fig. 4 d, f**, ***Supplementary Figure S9 b, d***). These findings suggest that each latent component represents features from distinct frequency bands, reflecting neural underpinnings of brain-symptom associations in a frequency- and symptom-specific manner. The contributions of node strengths to the latent components were generally stable, except for the theta band in the PS-derived components, but their contributions were minimal (***Supplementary Figure S7 b, Supplementary Figure S8 f***).

To confirm that brain-symptom associations were not due to sex or age effects, we performed the PLSC separately for males and females. The graph strength didn’t differ significantly between female and male cohorts neither for AC nor PS (two-tailed Mann-Whitney-U test, p>0.05) (***Supplementary Figure S10)***. Nevertheless, the latent brain-symptom associations exhibited distinct patterns for the female and male cohorts (***Supplementary Figure S11***). In the female cohort, beta-band (20–30 Hz) PS showed a significant correlation with depression severity (PHQ-9 and QIDS) and anxiety (GAD) (*p=0.0027*), whereas in the male cohort, depression severity (PHQ-9 and QIDS), well-being (WHO) and anhedonia (PVSS) were significantly associated with theta-alpha (4–9 Hz) band and beta-band (24–30 Hz) AC (*p=0.027*). The results thus suggest different brain-symptom association mechanisms for AC and PS coupling modes in males and females.

To assess age effects, we computed brain-symptom associations while regressing sex effects out. We obtained one significant latent component from AC (*p*=0.0145), while there were no significant latent components from PS (with the smallest *p* value being 0.0525) (***Supplementary Figure S12***). Contributions of symptoms and AC node strength to the latent components were, however, very similar to the results when both sex and age were regressed out, indicating that age did not substantially influence the results on AC.

For the HC cohort, we didn’t find any significant latent components with PLSC for AC (smallest *p*-value: 0.104) nor for PS (smallest *p*-value: 0.22), speaking against any significant brain-symptom associations.

### Depression phenotypes show distinct connectivity patterns and symptom profiles

Following the identification of brain-symptom associations in MDD, we applied the Leiden clustering method (Traag et al., 2019) to the two significant latent components to categorize MDD into distinct brain phenotypes. To determine the optimal resolution, we calculated modularity across a wide range of resolution values (**Fig. 5 a**). We obtained higher modularity values across the resolution range of 0.6–1.1, within which patients transitioned from being grouped into fewer and larger clusters to more numerous and smaller clusters (**Fig. 5 b**). This pattern reflects the emergence of increasingly fine-grained community structure as resolution increases. Using the elbow method to determine the optimal resolution (0.85), we obtained six distinct clusters with high modularity (**Fig. 5 c**). As one cluster contained only two patients, likely representing outliers, we excluded this cluster from further analyses. Including the second latent components from both AC and PS showed the robustness of clusters 2 and 3 but split the other clusters further (***Supplementary Figure S13***). Clusters were well separated by the two dimensions of brain-symptom AC and PS latent components (***Supplementary Figure S14***). Replication analysis, based on 1,000 iterations of the phenotyping pipeline using independent random subsamples that comprised 80% of the full cohort, demonstrated robust reproducibility and generalizability of the identified depression phenotypes (***Supplementary Figure S15***). Finally, we tested whether we could classify individuals to correct clusters using a decision tree classifier, and we found that the classification accuracy for each cluster could be as high as 98% (***Supplementary Figure S16***).

**Figure 5.**
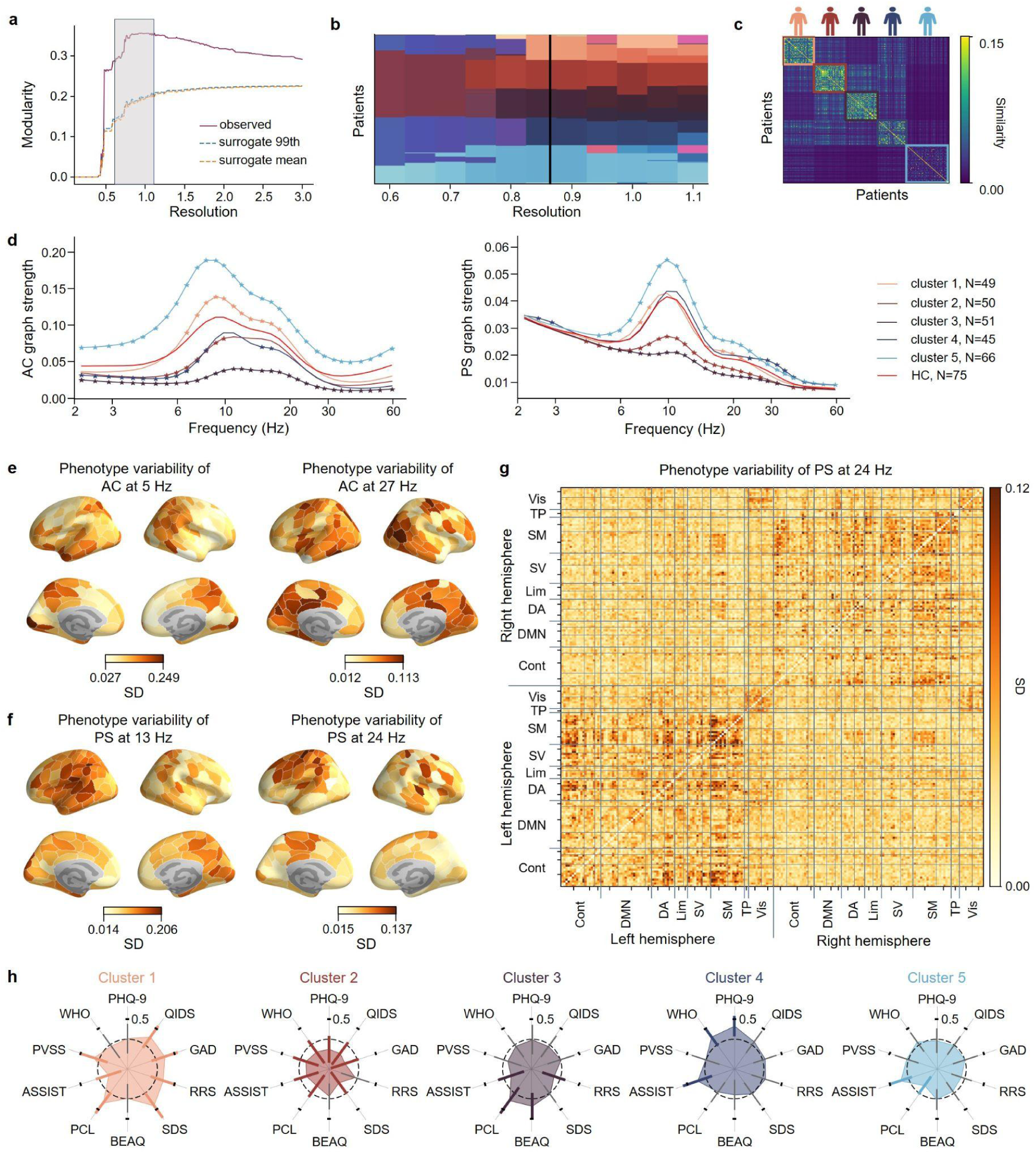
Low-dimensional brain-symptom associations delineate five depression phenotypes with distinct connectivity biomarkers and symptom profiles. **a** Modularity of observed data and surrogate data as a function of resolution in the Leiden clustering method. **b** Changes in clustering results for MDD patients as a function of resolution in the Leiden clustering method. **c** The similarity matrix between patients sorted by clusters. **d** Spectrum of graph strength of AC (left) and PS (right) for five biotypes and the HC group. Asterisks indicate statistical significance at p<0.05 (two-tailed Mann-Whitney U test, FDR-corrected with Benjamini-Hochberg method). **e–f** Standard deviations of node strength across clusters for AC at 5 and 27 Hz (**e**) and for PS at 13 and 24 Hz (**f**). Averaged node strength for each cluster was min-max normalized to the range 0–1, and then their standard deviations were calculated across clusters for each parcel. **g** Phenotype variability of PS connectivity strength between parcels at 24 Hz. Averaged connectivity strength for each cluster was min-max normalized to the range of 0–1, and then their standard deviations were calculated across clusters. **h** Symptom profiles (z-scored) for the five phenotypes. The dashed circle at a value of 0 represents the mean of z-scored symptom scales across the whole clinical cohort. The colored and thick color bars in each cluster represent statistically significant differences compared to the other clusters (bootstrapping, N=10,000, at p<0.05).

Having established the robustness of the phenotypes, we subsequently examined whether PS and AC for each cluster differed significantly from the HC group. At both whole-brain (**Fig. 5 d**) and parcel levels (***Supplementary Figure S17***), all five depression phenotypes demonstrated distinct spectral profiles differing from the HC group for both PS and AC in a frequency-specific manner. Cluster 1 exhibited alpha-to-beta-band hyperconnectivity in AC and PS, with a wider frequency band for AC (7–20 Hz) than PS (7–9 Hz and 18–22 Hz). Similarly, cluster 5 exhibited overall robust hyperconnectivity both in AC (2–60 Hz) and PS (6–30 Hz). Clusters 2–4 exhibited, in contrast, widespread hypoconnectivity in a frequency-specific manner. Cluster 2 exhibited hypoconnectivity, particularly in AC at the theta-alpha band (5–10 Hz) and in PS within the alpha band (8–15 Hz), cluster 3 over a wide frequency range in AC (2–60 Hz) and in PS (6–30 Hz), and cluster 4 in AC within the delta-theta range (2–8 Hz) but hyperconnectivity in PS in the beta-gamma band (25–40 Hz). Interestingly, graph strength in the alpha band peaked at different frequencies across clusters for AC, specifically at 9, 11, 11, 10, and 8 Hz for clusters 1–5, respectively, whereas PS showed much more consistent peak frequencies, with peaks at 10, 10, 11, 10, and 10 Hz for clusters 1–5 (**Fig. 5 d**). For the oscillatory networks that contributed most strongly to brain-symptom associations (**Fig. 4 d**), we observed phenotype variability in anatomical patterns across clusters. Specifically, AC at 5 Hz showed greater variability across clusters in the posterior parietal cortex (PPC) and early visual cortices, while AC at 27 Hz varied more across clusters in somatomotor (SM) areas and widely over visual areas and posterior cingulate cortices (**Fig. 5 e, *Supplementary Figure S18 a–b***). In contrast, PS at 13 Hz showed higher variability across clusters in the ventral prefrontal cortex (PFC), SM, PPC, and areas of the limbic network including the posterior cingulate, while PS at 24 Hz varied more across PFC and SM areas (**Fig. 5 f, *Supplementary Figure S18 c–d***). At the edge level, AC at 5 Hz reflected phenotypic variability in connectivity strength both within and between hemispheres (***Supplementary Figure S19 a***), and AC at 27 Hz showed greater phenotypic variability within the visual network and between the visual and default mode networks (***Supplementary Figure S19 b***). PS at 13 and 24 Hz exhibited increased phenotypic variability within each hemisphere (**Fig. 5 g, *Supplementary Figure S19 c–d***). These findings highlight the spectral variability that differentiates the depression phenotypes.

No significant differences between clusters were found in gender (***Supplementary Figure S20 a***, Chi-square test at *p*>0.05, FDR corrected) and age (***Supplementary Figure S20 b***, two-tailed Mann-Whitney U test at *p*>0.05, FDR corrected) distributions. We then tested whether symptom profiles of one phenotype differed significantly (bootstrapping, *N*=1,000, *p*<0.05) from the rest of the cohort (**Fig. 5 h**). All symptom scores were z-scored across the whole cohort, so that the zero circle represented the average scores after normalization. WHO and PVSS scores were inverted (multiplied with –1) before z-scoring to maintain the direction of meaning, ensuring that a higher normalized score consistently indicated more severe symptoms. The values outside the zero-circle indicated more severe symptoms, while values below 0 indicated less severe symptoms. Cluster 1, associated with hyperconnectivity, exhibited more severe symptoms for most questionnaires, namely QIDS, GAD, RRS, SDS, PCL, and PVSS. Cluster 2, exhibiting hypoconnectivity in both AC and PS, demonstrated generally less severe symptomatology compared to the other depression phenotypes except for RRS and BEAQ. Cluster 3, with more robust hypoconnectivity, had larger PCL and BEAQ scores, while RRS and ASSIST scores were smaller. Cluster 4, with AC hypoconnectivity, was characterized by higher PHQ-9, ASSIST, and WHO scores. Cluster 5, with the highest hyperconnectivity in AC and PS, was characterized by more severe ASSIST but smaller PCL scores. Overall, these data established that the identified MEG oscillation phenotypes were characterized by different symptom profiles, representing different dimensions of depression symptoms.

## Discussion

### Depression phenotypes show distinct oscillatory connectivity patterns

MDD is a multifactorial disorder characterized by symptom heterogeneity and complex comorbidities. Previous evidence supports the existence of distinct depression phenotypes, based on genetic, neuroimaging, biomedical, and psychometric measurements (Beijers et al., 2019). While studies using fMRI-based functional connectivity have advanced our understanding of the biological basis of symptom-specific depression heterogeneity (Drysdale et al., 2017; Dunlop et al., 2024), such biotypes have not been identified for electrophysiological connectivity data, despite the fact that they reflect the underlying biological pathophysiologies.

Here, we demonstrate for the first time that individually source-reconstructed MEG data and oscillation-based connectomics can identify biological depression phenotypes. With this approach, we identified five stable depression phenotypes showing neurophysiological and symptomatological heterogeneity. These phenotypes exhibited frequency-specific hyperconnectivity or hypoconnectivity compared to the healthy controls. Such increases and decreases in oscillation-based connectivity indicate separate pathophysiological mechanisms linked to depression across the affected population. The presence of opposing phenotypes in depression might explain the discrepancies in the previous findings of both increased (Miljevic et al., 2023; Saletu et al., 2010; Whitton et al., 2018) and decreased alpha connectivity (Nugent et al., 2020) as well as the recent negative findings of the meta-analysis of fMRI data (Winter et al., 2022).

Critically, oscillations are one of the intermediate stages linking cellular and molecular deficiencies to the whole-brain level and to behavior (Varela et al., 2001) mechanism to explain the complex relationship between treatment modalities in depression (Leuchter et al., 2015). Our results of altered large-scale oscillatory networks in depression are in line with earlier findings of inhibitory deficiencies, specifically in GABAergic STT-interneurons in cellular models of depression, suggesting that inhibitory dysfunctions may contribute to network-level aberrancies that cause symptoms at the behavioral level (Fee et al., 2017; Fogaça & Duman, 2019; Lin & Sibille, 2015).

### Oscillation Phenotypes are associated with different symptom profiles

Although biological phenotypes are meaningful per se for understanding the pathophysiological basis of heterogeneity in depressive symptoms, the critical question is whether the phenotypes are clinically relevant to understand the heterogeneity of depressive symptoms. Our results show that the five MEG connectivity phenotypes exhibited statistically different symptom profiles. One phenotype (cluster 1) exhibiting moderate theta-beta band hyperconnectivity was characterized by more severe symptoms compared to the other phenotypes across most domains, including higher depressive symptoms, anxiety, rumination, functional impairment, PTSD symptoms, and anhedonia, and thus represented the main depression phenotype. In contrast, another phenotype (cluster 5) with severe hyperconnectivity across the whole frequency spectrum differed from other phenotypes only by more severe substance abuse and weaker PTSD symptoms, thus being a specific phenotype for addiction dominant disorder. This suggests that while these symptom characteristics may both relate to hyperconnectivity, they emerge via different biological mechanisms.

In line with the oscillation connectivity phenotypes being linked to the symptomatology, the phenotypes with hypoconnectivity also showed specific symptom profiles. The dissociation of the PTSD phenotype (cluster 3) from the more typical MDD phenotypes is unique and highlights the specificity of these symptoms. The result aligns with the recent identification of distinct molecular pathologies and markers for PTSD and MDD using brain multi-omic molecular dysregulations (Daskalakis et al., 2024).

### Oscillatory phenotypes are associated with distinct spatial patterns and alpha peak frequency

The hypo- and hyperconnectivity maps across frequencies and phenotypes were spatially distributed. Specifically, the theta-band AC and alpha-band PS connectivity showed phenotypic variability in parcels of the PFC which was recently shown to be expanded in individuals with depression (Lynch et al., 2024). In addition, the findings of limbic and cingulate connectivity abnormalities are in line with the fMRI-based connectivity phenotypes (Drysdale et al., 2017; Dunlop et al., 2024) and with findings of negative self-referential processing in MDD (Sheline, Barch, Price, Rundle, Vaishnavi, & Snyder, 2009; Sheline et al., 2010; Wise et al., 2017). The distributed oscillation connectivity patterns further speak in favor of the idea that depression would especially be due to the disruption of brain dynamics and changes in network interactions (Piguet et al., 2021). Our findings on correlations between theta hyperconnectivity with ruminative and PTSD symptoms aligns well with evidence that excessive low-frequency network connectivity underlies self-focused thought and negative affect (Dell’Acqua et al., 2020) and ruminative symptoms (Benschop et al., 2021). Phenotypic variability at beta band for both PS and AC connectivity were found in the sensorimotor areas, possibly reflecting psychomotor symptoms and sensorimotor slowing down (Sobin & Sackeim, 1997), which have been linked to hyperconnectivity in the thalamo-cortical network that includes SM areas (Wüthrich et al., 2024).

Crucially, we found that oscillation connectivity phenotypes were associated with variations of individual alpha peak frequency (iAF) of AC. The iAF has been implicated in various mental disorders and is considered as a neurophysiological marker of cognitive functioning. Higher iAF has been associated with enhanced cognitive performance, whereas lower iAF has been linked to increased symptom severity across multiple psychiatric conditions (Arns et al., 2010; Grandy et al., 2013; Voetterl et al., 2023). Consistent with these findings, cluster 1, characterized by a lower iAF, was associated with more severe depressive symptoms, while cluster 2 with higher iAF was associated with less severe symptoms. These findings support the potential utility of iAF as a biomarker for stratifying depression phenotypes (Voetterl et al., 2023).

The incidence and symptoms of depression have been shown to vary across age (Collier Villaume et al., 2023; Dunlop et al., 2021; Fjell & Walhovd, 2010; Hybels et al., 2012) and gender (Joel et al., 2015; Piccinelli & Wilkinson, 2000; Salk et al., 2017; Talishinsky et al., 2022; Valentino, 2024). We found gender differences in brain-symptom associations, which suggest that partially distinct oscillatory coupling mechanisms may underlie depressive symptoms in males and females. These differences may reflect variations in hormonal influences, neuroplasticity, and network-level connectivity dynamics, contributing to sex-specific neurophysiological mechanisms in depression (Bangasser & Cuarenta, 2021; Jaworska et al., 2012; Kuehner, 2017; Labaka et al., 2018). Our findings highlight the importance of considering gender-specific functional connectivity alterations in depression, which could inform more personalized interventions.

### Conclusions

Intervention outcomes for depression show large heterogeneity across patients (Liao et al., 2025; Serretti, 2025; C. Wu et al., 2025). Here we identified, in a cohort of 263 MDD patients, novel phenotypes based on oscillatory connectivity that were distinct both in spectral patterns and symptom profiles. Given that oscillations are among the key mediators between cellular neurophysiology and behavior, these phenotypes may pave the way for accurate intervention outcome prediction models and uncover novel mechanistic biomarkers.

## Methods

### Recruitment

These data were collected as a substudy of the clinical trial “Meliora RCT”, a remote, randomized, double-blinded, comparator-controlled, cross-over, add-on, and three-arm clinical device trial with 18–65-year-old adults with an interview-confirmed MDD diagnosis. The study was approved by the Helsinki University Hospital Regional Committee on Medical Research Ethics (HUS/3042/2021) and the Finnish Medicines Agency (FIMEA/2022/002976) and conducted in compliance with the Declaration of Helsinki. The main study was pre-registered on ClinicalTrials.gov (NCT05426265). All patients gave their informed written consent prior to participation.

The study patients were recruited in collaboration with Finnish healthcare partners (Helsinki University Hospital, Turku University Hospital, and Mehiläinen Ltd.), with whom research permits were signed and where clinicians were encouraged to share information about the study to their patients. Participants were also recruited through social media (Facebook, Instagram, and Reddit), email campaigns, and posters placed at university campuses. All recruitment channels guided the interested patients to the study website, where they digitally signed an informed consent form.

### Participants

The study patients were adults aged 18–65 years with interview-confirmed major depressive disorder (MDD). Patient eligibility was evaluated in a phone interview conducted by Clinical Subject Coordinators (CSC). The CSCs evaluated MDD using the Mini-International Neuropsychiatric Interview (M.I.N.I.) 6.0.0. module A (Sheehan et al. 1998), which is based on the Diagnostic and Statistical Manual of Mental Disorders 4th edition criteria (DSM-IV). The patients were required to have an existing mental health treatment contact. Patients with suicidality were excluded from the study. When necessary, the CSCs evaluated suicidality using the MINI module B, where a score of 17 or more was considered an absolute exclusion criterion. Patients with severe self-report gaming addiction were also excluded. When necessary, the CSCs evaluated patient gambling addiction with the Canadian Problem Gambling Index (PGSI) (Ferris & Wynne, 2001), where a score of 8 or more served as exclusion criteria, and digital gaming addiction with a shortened 7-item version of the Gaming Addiction Scale (GAS-7) (Lemmens, Valkenburg & Peter, 2009), where 4 or more answers of “sometimes”, “often” or “very often” constituted exclusion criteria. Patients who were unable to consent, pregnant or nursing, inmates or forensic patients, had a psychotic disorder, or had a neurological disorder such as epilepsy or brain injury (migraine didn’t prevent participation), were excluded.

Study participants consisted of the volunteers from the main study with *N*=263 patients with major depressive disorder (163 females, 87 males, and 13 labeled as “other”) and 75 healthy control participants (29 females, 45 males, and 1 labeled as “other”). Criteria for study inclusion for both groups included age in the range 18-65, normal or corrected to normal vision, and compatibility with neuroimaging methods.

### Clinical diagnosis

To assess the acuteness and severity of MDD symptoms, a brief structured diagnostic interview, the M.I.N.I. module (Sheehan et al., 1998), was conducted during the phone interview. Requirement for study inclusion for the MDD patients was a score of 5 or more and a “yes” answer to question A4 (“Have the symptoms caused you significant problems at home, school, in social, professional or other important areas of activity?”). For healthy control participants, the requirement for study inclusion was a score of less than 5 in the assessment. Interviewers were experienced health care professionals or students educated in the use of the M.I.N.I. interview. If the interviewee expressed suicidal ideation, the interviewer asked for clarification questions and if, according to the estimation of the interviewer, there was concern for risk of self-harm, the interviewee could not be included into the study and was encouraged to reach out for their health care provider. If the interviewer was uncertain of the severity of suicidality, the M.I.N.I. module B (Suicidality) could be conducted, in which a score of 17 or more was a definite indication for exclusion. In addition, the interviewer could, at their discretion, exclude the interviewee with a lower score.

### Symptom measures

All participants were required to complete 10 following self-report symptom scales to assess various aspects of mental health and functioning. These scales include Patient Health Questionnaire (PHQ-9) measuring depressive symptoms (range: 0–27), Quick Inventory of Depressive Symptomatology 16-item (QIDS) evaluating depressive symptom severity (range: 0–27), Generalized Anxiety Disorder 7-item (GAD) Scale assessing anxiety symptoms (range: 0–21), Ruminative Responses Scale Short-version (RRS) evaluating ruminative thought patterns (range: 8–32), Sheehan Disability Scale (SDS) evaluating functional impairment (range: 0–30), Brief Experiential Avoidance Questionnaire (BEAQ) measuring experiential avoidance (range: 15–90), PTSD Checklist for DSM-5 (PCL) assessing post-traumatic stress disorder symptoms (range: 0–80), Alcohol, Smoking, and Substance Involvement Screening Test Lite (ASSIST) assessing substance use involvement (range: 0–20), Positive Valence Systems Scale (PVSS) examining positive affect and reward processing (range: 21–189), and WHO-5 Well-Being Index (WHO) measuring subjective well-being (range: 0–25). Total PHQ-9 scores of 5, 10, 15, 20 represent cut points for mild, moderate, moderately severe and severe depression, respectively. For evaluation, symptom scores were converted to either a min-max range from 0 to 1, or z-scored. In either case, PVSS and WHO were inverted, so that for them, as for the other scores, a higher value represented more severe symptoms (lower positive valence or greater anhedonia for PVSS and worse well-being for WHO).

### Neuroimaging data acquisition

Fifteen minutes eyes-open resting-state MEG data were recorded with a 306-channel MEG system (TRIUX or TRIUXneo, MEGIN Oy, Helsinki, Finland; 204 planar gradiometers and 102 magnetometers) at the BioMag Laboratory, HUS Medical Imaging Center, Helsinki, Finland or at MEG Core, Aalto University, Espoo, Finland. Bipolar horizontal and vertical electrooculography (EOG) and electrocardiography (ECG) were recorded for detection of eye movements and cardiac artefacts. Participants were instructed to sit in a dimly lit room and to focus on a fixation cross. T1-weighted anatomical MRI scans were obtained with a 3-tesla whole-body MRI scanner (Magnetom Skyra, Siemens, Erlangen, Germany) at AMI Centre, Aalto University at a resolution of 0.8×0.8×0.8 mm, repetition time of 2530 ms, and echo time of 3.42 ms.

### MEG data preprocessing and source modeling

Temporal signal space separation (tSSS) in the Maxfilter software (Elekta Neuromag) was used to suppress extracranial noise from MEG sensors, interpolate bad channels, and compensate for head motions. 50-Hz line noise and its harmonics were removed with a Finite Impulse Response (FIR) notch filter. Independent components analysis was used to remove ocular, heartbeat, and muscle artifacts. We used the FreeSurfer software (https://surfer.nmr.mgh.harvard.edu/) for volumetric segmentation of MRI data, surface reconstruction, flattening, and cortical parcellation. Source reconstruction was performed with minimum norm estimation (MNE) using dynamic statistical parametric maps (dSPM) with the MNE software (https://mne.tools/stable/index.html). A surface-based source space with 5-mm spacing and 1-layer (inner skull) symmetric boundary element method (BEM) was used in computing the forward operator. Noise covariance matrices (NCM) were obtained from preprocessed data filtered to 151–249 Hz. We then estimated vertex fidelity by applying forward and inverse operators to complex white-noise time series and computing the correlation between original and forward-inverse-modeled time series (Korhonen et al., 2014). The obtained fidelity-weighted inverse operators were used for collapsing vertex time series into the 200 parcels of the Schaefer atlas (Schaefer et al., 2018). Broadband parcel time series were then filtered into narrowband time series with 32 Morlet wavelets with center frequencies spanning from 2.1 to 59.3 Hz in log-linear space.

### Phase synchrony and amplitude correlation analysis and statistical analysis

Phase synchrony (PS) and amplitude correlations (AC) are two intrinsic modes of functional connectivity in electrophysiological data, which are believed to represent complementary, although related, mechanisms of neuronal communication (Engel et al., 2013; Siebenhühner et al., 2024; Siems & Siegel, 2020; Zhigalov et al., 2017). We applied weighted phase lag index (wPLI) (Vinck et al., 2011) to assess PS and orthogonalized correlation coefficient (oCC) to measure AC which are maximally insensitive to false-positive interactions due to the source leakage (J. M. Palva et al., 2018).

Phase synchrony was computed between the two time series *x*(*t*) and *y*(*t*) with wPLI (Vinck et al., 2011) as:

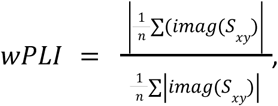

where *S*_*xy*_ is the cross-spectrum of *x*(*t*) and *y*(*t*) and *imag* is the imaginary part.

Amplitude correlations were computed between the amplitude envelopes of the narrowband time series *x*(*t*) and *y*(*t*) with the orthogonalized correlation coefficient (oCC), where the time series *x*(*t*) is orthogonalized with respect to *y*(*t*) (Brookes et al., 2012; Hipp et al., 2012).

AC and PS were computed between all pairs of parcels and for each Morlet Wavelet frequency. The resulting adjacency matrices were represented as weighted graphs (Bullmore & Sporns, 2009). As a measure of global connectivity, we used graph strength (GS) that was computed as the mean connectivity between all parcel pairs. As a measure of local connectivity, we used node strength (NS) that was computed by averaging the connectivity strength (wPLI or oCC) of a given parcel with all other parcels.

To reduce the risk of conflating general demographic trends, we regressed out the effects of age and gender for PS and AC node strength values with multivariate linear regression.

### Statistical analysis between the cohorts

Statistical analyses were performed to compare PS and AC between the MDD and HC cohorts both at the graph and parcel levels. At the graph level, the graph strength for each frequency was compared between two cohorts using a two-tailed Mann-Whitney U test (p<0.05). Multiple comparisons were corrected separately for PS and AC using the False Discovery Rate (FDR) method. At the parcel level, node strength for each node and each frequency was similarly compared using a two-tailed Mann-Whitney U test (p<0.05) with FDR correction applied separately for PS and AC.

### Brain-symptom associations and statistical analysis

Univariate correlations between PS/AC node strength and each symptom score were computed using the Pearson correlation method separately for each parcel and frequency. For each symptom, we then calculated the fraction of parcels showing statistically significant correlations for each frequency. To control multiple comparisons, we performed permutation-based statistics to obtain null distributions with 1,000 randomizations by randomly shuffling symptom scores across patients and keeping node strength unchanged. We then derived the 97.5th percentile thresholds of the fraction of significant parcels for both positive and negative correlations and subtracted the remaining expected false positive rate from the observed fraction of significant parcels (Puoliväli et al., 2020).

Multivariate brain-symptom correlations were obtained with Partial Least Squares Correlation (PLSC) (Krishnan et al., 2011; McIntosh & Lobaugh, 2004; Vieira et al., 2024). PLSC aims to extract latent components which can maximize the common information shared between two datasets by projecting raw data into a low-dimensional space.

For two given matrices *X* and *Y*, where *X* contains vectorized connectivity data from *N* subjects and *Y* contains the behavioral data from the same subjects, singular value decomposition (SVD) is applied to the cross-covariance matrix *R* between *X* and *Y*, resulting in 3 low-dimensional matrices: *U*, *S* and *V*:

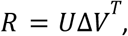

where *R* = *Y*^T^*X*. Note that *R* is also a Pearson’s correlation matrix, because *X* and *Y* are expressed as z-scores. *U* and *V* are behavioral and imaging saliences which represent the contribution of the raw features on the latent components. Then the latent components are computed by projecting the raw matrices onto the saliences:

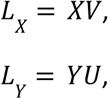

Where *L*_*X*_ and *L_Y_* are called brain scores and behavior scores, respectively. Imaging and behavioral structure coefficients (or “loadings”), *corr*(*X*, *L_X_*) and *corr*(*Y*, *L_Y_*) are computed by Pearson’s correlations. Compared with saliences, loadings are recommended to present the contributions of raw features to latent components due to the multicollinearity of data and easy interpretation (values are bounded between –1 and 1).

To examine how the fixed effect model can be extended to a random effect model which applies to the broader population, we followed well-established approaches (Krishnan et al., 2011; McIntosh & Lobaugh, 2004). Statistical significance of the latent components was obtained using a permutation test by randomly shuffling the order of *X* 1,000 times and building the null distributions of singular values. To test the generalizability and stability of loadings, we created 1,000 bootstrap samples by sampling with replacement and estimated the standard errors for each element’s loading of each latent variable from bootstrap samples. As the ratios of loadings and the corresponding standard errors are akin to a z-score, the loadings where absolute values of ratios were larger than 2 were significantly stable. A technical problem with permutations and bootstrapping is that resampling may cause axis rotation and reflection, which can make a random effect model not comparable to the fixed effect model. We applied Procrustes rotation to correct the rotations and reflections in saliences (Krishnan et al., 2011; McIntosh & Lobaugh, 2004).

### Identification of phenotypes and statistical analysis

We used significant latent components to identify phenotypes with the Leiden community detection algorithm, designed to identify non-overlapping communities from a network (Traag et al., 2019). A key feature of the Leiden method is the resolution parameter that determines the scale of community detection, yielding a balance between finer, smaller clusters and coarser, larger ones. This adaptability enables the clustering granularity to align with the structural complexity of the data.

The patient-patient Euclidean distances (dissimilarity matrix) were computed with the selected latent components and then transferred to the similarity matrix using the transformation 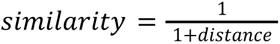, which was used as input for Leiden clustering. We performed Leiden clustering at multiple resolutions, obtaining a partition for each resolution, and calculated the modularity as a function of resolution.

To assess the statistical significance of each partition, we implemented a permutation-based significance test to generate a null distribution of modularity for each resolution by randomly permuting the assignment of patients to clusters for 1,000 times and derived the mean and the 99th percentile modularity values from this null distribution. The elbow method was used to determine the optimal partitions based on significant modularity values.

Given the sample size of our study, we also performed an in-sample replication analysis to evaluate the reproducibility and generalizability of the identified depression phenotypes. We repeated the entire phenotyping pipeline, from brain-symptom association analysis to clustering, 1,000 times, each time using a randomly selected 80% subset of the full clinical cohort. To assess cluster stability, we then calculated the probability that a given pair of patients were assigned to the same cluster across the iterations in which both patients were included.

### Characterization of phenotypes and statistical analysis

To assess how connectivity in each phenotype deviates from that in the HC group, we performed statistical testing on graph strength and node strength between each depression phenotype and the HC group for each frequency. Pairwise statistical analyses were conducted using the two-tailed Mann-Whitney U test at *p*<0.05, corrected for multiple comparisons using the Benjamini-Hochberg procedure. Additionally, we assessed differences in the two significant brain scores between each pair of depression phenotypes with the two-tailed Mann-Whitney U test at *p*<0.05, corrected with the Benjamini-Hochberg method for multiple comparison correction.

For comparing and visualizing the symptom profiles of the discovered phenotypes, all symptom scores were z-scored across the whole clinical cohort, so that the zero circles on the spider plots represents the mean of all patients. For each symptom scale, we assessed the statistical differences between patients in one phenotype against others with a bootstrap resampling procedure (*N*=1,000). We estimated the bootstrap mean value and the confidence intervals at the alpha level of 0.05 (confidence limits from 2.5th–97.5th percentiles) for one phenotype and the others. If the bootstrap mean value of the other phenotypes are out of the range of confidence intervals of this phenotype, the differences will be considered as significant. Age and gender differences between pairs of clusters were evaluated using the two-tailed Mann-Whitney U test and the Chi-square test, respectively, with a significance threshold of *p*<0.05. To account for multiple comparisons, the Benjamini-Hochberg procedure was applied to control the false discovery rate.

## Data availability

The raw datasets generated and analyzed in the current study are not publicly available due to them containing information that could compromise research participant privacy or consent. A minimal dataset that can be used to reproduce main findings of this study will be uploaded to Datadryad.

## Code availability

MRI data were processed using FreeSurfer software (v.7.3.2) (https://surfer.nmr.mgh.harvard.edu/). MEG preprocessing and source reconstruction was done with the MNE-python package (https://mne.tools/stable/index.html). Code for computing phase synchrony and amplitude correlations can be found at (https://github.com/palvalab). Code for conducting PLSC analysis was customized from the myPLS toolbox (https://github.com/MIPLabCH/myPLS). For the Leiden clustering method, we used the leidenalg package (https://github.com/vtraag/leidenalg).

## Acknowledgements

This work was supported by Ella and Georg Ehrnrooth Foundation to W.L., and Finnish Cultural Foundation grants 00220945 and 00242647 to F.S. Work on “PlaStim: Plasticity Stimulation in the Treatment of Anhedonia” was supported by Wellcome Leap as part of the Multi-Channel Psych Program to S.P. and J.M.P.

## Author contributions

W.L. conceived the study, preprocessed MEG and MRI data, developed the methods, performed data analysis, and drafted the manuscript. M.V. recruited subjects and collected MEG data. P.P recruited subjects and collected MRI data. A.A. collected MEG data and preprocessed MEG data. S.K. processed MRI data. J.J. contributed to the MEG preprocessing pipeline, data analysis pipeline, and data visualization. F.S. preprocessed MEG data and drafted the manuscript. A.S. collected symptom data. H.R. contributed to ethical application of the study. R.I. acquired funding. E.C. acquired funding. E.I. contributed to the clinical investigation and study design. D.V.D.V. contributed to data analysis pipeline design. J.M.P conceived the study and acquired funding. S.P. conceived and supervised the study, acquired funding, and drafted the manuscript. All authors contributed to writing the manuscript.

## Competing interests

The authors declare no competing interests.

## Notes

### Competing Interest Statement

The authors have declared no competing interest.

