## Supplementary figures for "Low-dimensional brain-symptom associations delineate depression phenotypes with distinct connectivity biomarkers and symptom profiles"

### Supplementary Figure S1

Age distribution for both clinical and control cohorts and comorbidity of the clinical cohort. **a** The distribution of age for each gender category for the major depressive disorder (MDD) cohort. No significant differences were found between any pair of gender categories with the Mann-Whitney U test at the level of 0.05. (Mann-Whitney U test, $p$=0.061 for male-female, $p$=0.536 for female-others, and $p$=0.155 for male-others). **b** The distribution of age for each gender category for the healthy control (HC) cohort. No significant differences in age were found between the male and female cohorts (Mann-Whitney U test, $p$=0.458). **c** Percentages of the 13 most common comorbidity combinations (including patients who were only diagnosed with depression), collectively accounting for 82.13% of all patients. 32.70% of patients were diagnosed with depression alone, 21.30% were diagnosed with both anxiety and depression, and 6.84% were diagnosed with ADHD in addition to depression.

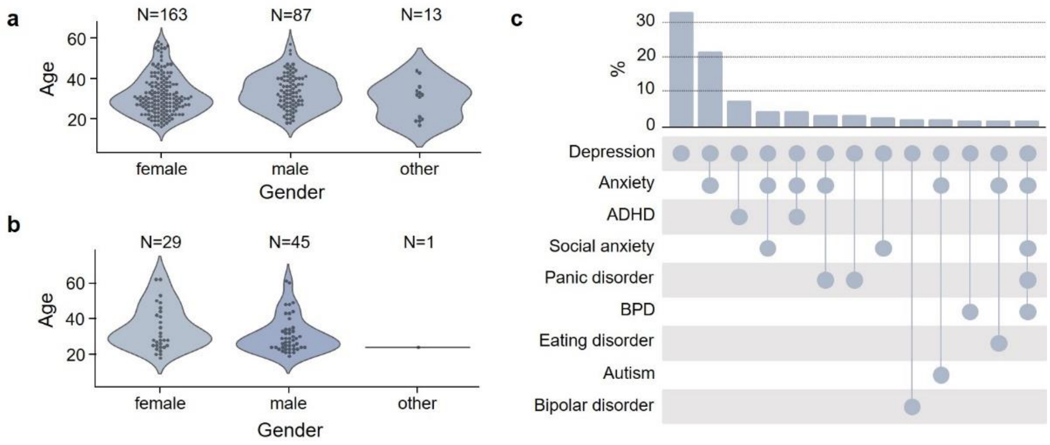

#### Supplementary Figure S2

Symptom profiles for females (left) and males (right) for the MDD cohort. All symptom scores were z-scored across the whole MDD cohort. The dashed circle, representing the value of zero, is the mean of z-scored symptom scores across the whole MDD cohort. The colored and thick color bars represent the statistical significance of differences compared between the male and female cohorts with 1,000 bootstrapping at the level of 0.05. Error bars represent the confidence intervals of the bootstrapping mean value (confidence limits from 2.5%–97.5%).

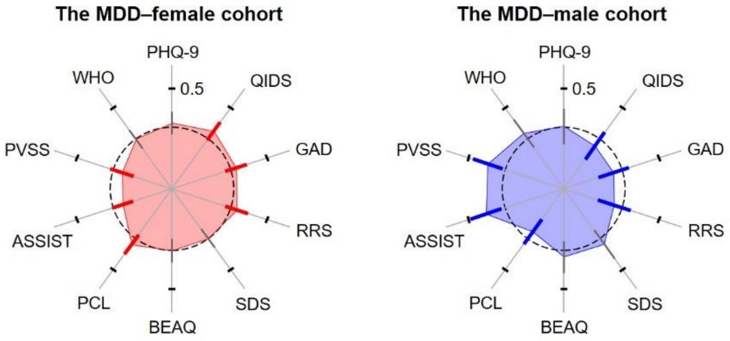

### Supplementary Figure S3

The spectrum of amplitude for the MDD and HC cohorts. **a** The confidence intervals (CIs) were obtained by the 2.5th and 97.5th percentiles on the mean amplitude from 1,000 bootstrapping. **b** The CIs were obtained by the 2.5th and 97.5th percentiles on amplitude across subjects. No significant differences in amplitude were found between MDD and HC cohorts (Mann-Whitney U test, *p*>0.05, False Discovery Rate (FDR) corrected with the Benjamini-Hochberg procedure).

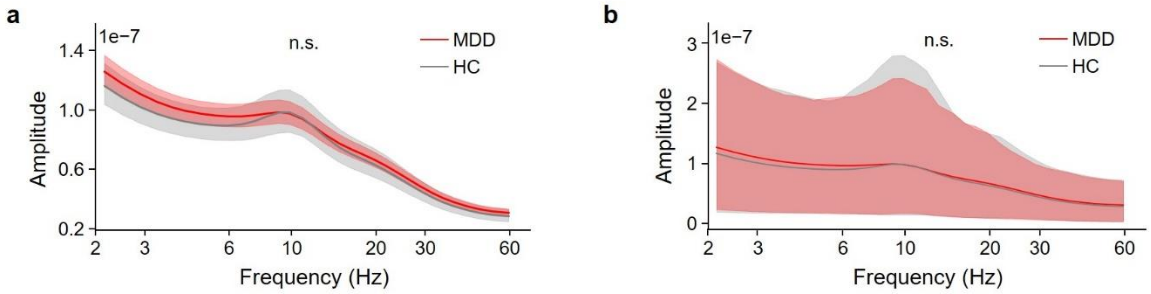

#### Supplementary Figure S4

The spectrum of graph strength of amplitude correlations (AC) (**a**) and phase synchrony (PS) (**b**) for the MDD and HC cohorts. The CIs were obtained by the 2.5th and 97.5th percentiles across subjects. No significant differences for both AC and PS graph strength at each frequency were found between MDD and HC cohorts (Mann-Whitney U test, $p$>0.05, False Discovery Rate (FDR) corrected with the Benjamini-Hochberg procedure).

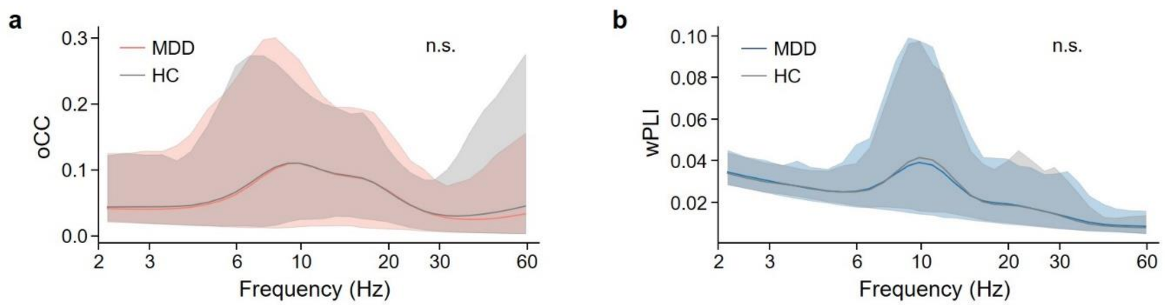

### Supplementary Figure S5

**a** The contrast of AC node strength between the MDD and HC cohorts at 2.15, 2.49, 2.88, and 3.31 Hz on parcels where differences were significant between two cohorts (Figure 3**C**). **b** The contrast of PS node strength between the MDD and HC cohorts at 2.88, 4.15, 10.92, and 19.7 Hz on parcels where differences were significant between two cohorts (Figure 3**D**).

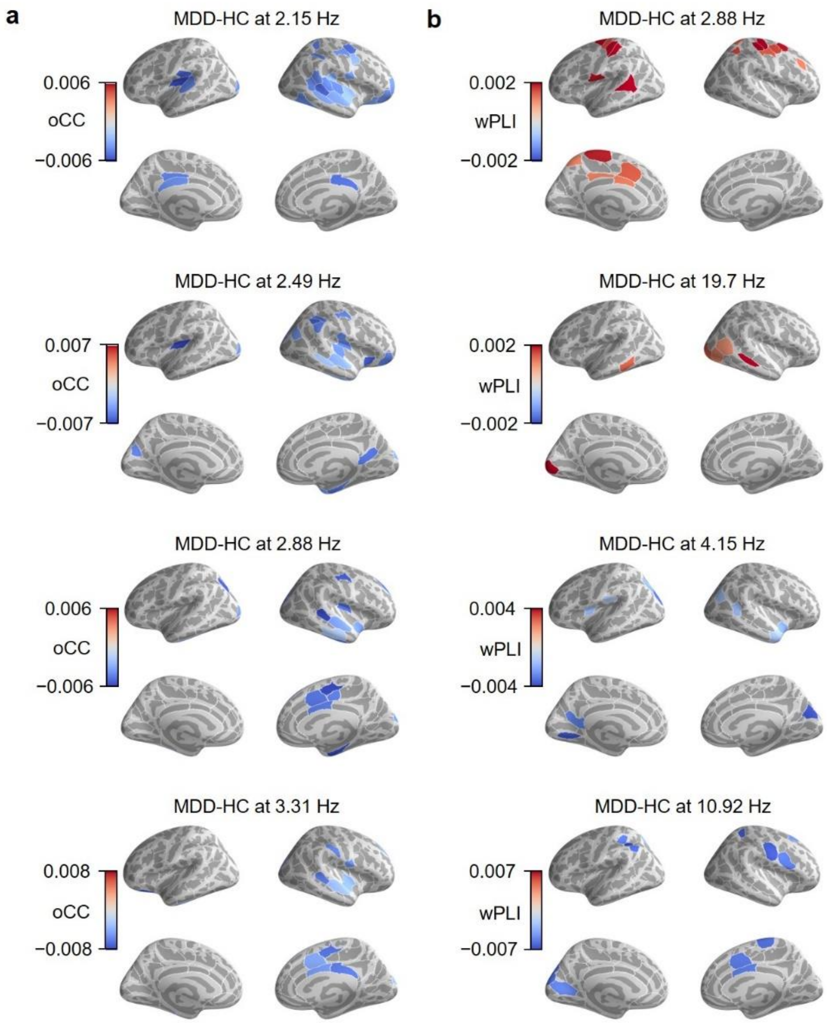

### Supplementary Figure S6

Spearman correlations between AC and PS graph strength for both MDD and HC cohorts. Asterisk represents significant correlations (Spearman's *r*, FDR-corrected with Benjamini-Hochberg procedure at $p<0.05$).

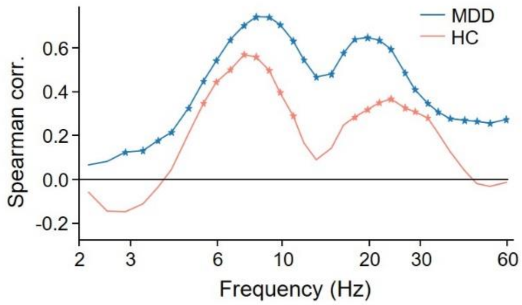

### Supplementary Figure S7

Stability of symptom and node strength contributions to significant latent components (one from AC and one from PS). **a** Stability of symptom contributions to the significant components from AC and PS. If the stability index is larger than 2 or smaller than -2, then the contribution was considered as stable. **b** The fraction of parcels that their node strength has stable contributions to the significant latent components from AC and PS.

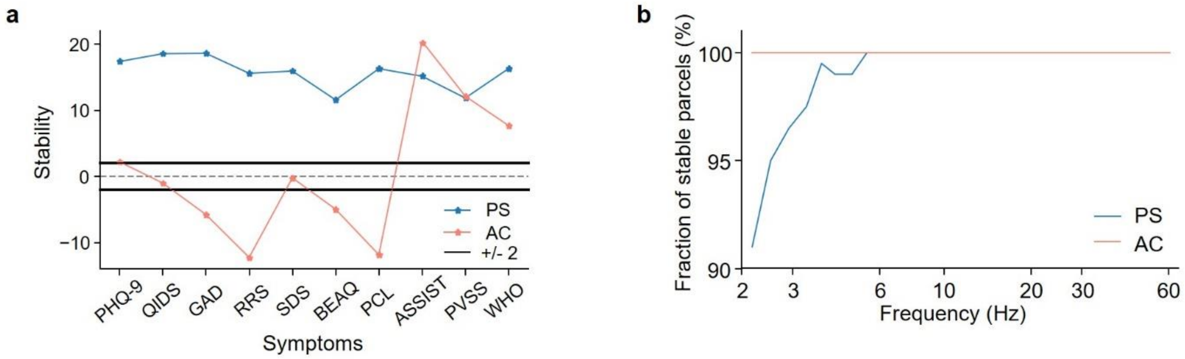

### Supplementary Figure S8

Contributions of AC and PS on latent brain-behavior associations. Fraction of parcels per frequency among the top 5% (**a**), 10% (**b**), 20% (**c**), 30% (**d**), 40% (**e**), and 50% (**f**) of AC or PS node strength that contributed the most to the two significant latent components.

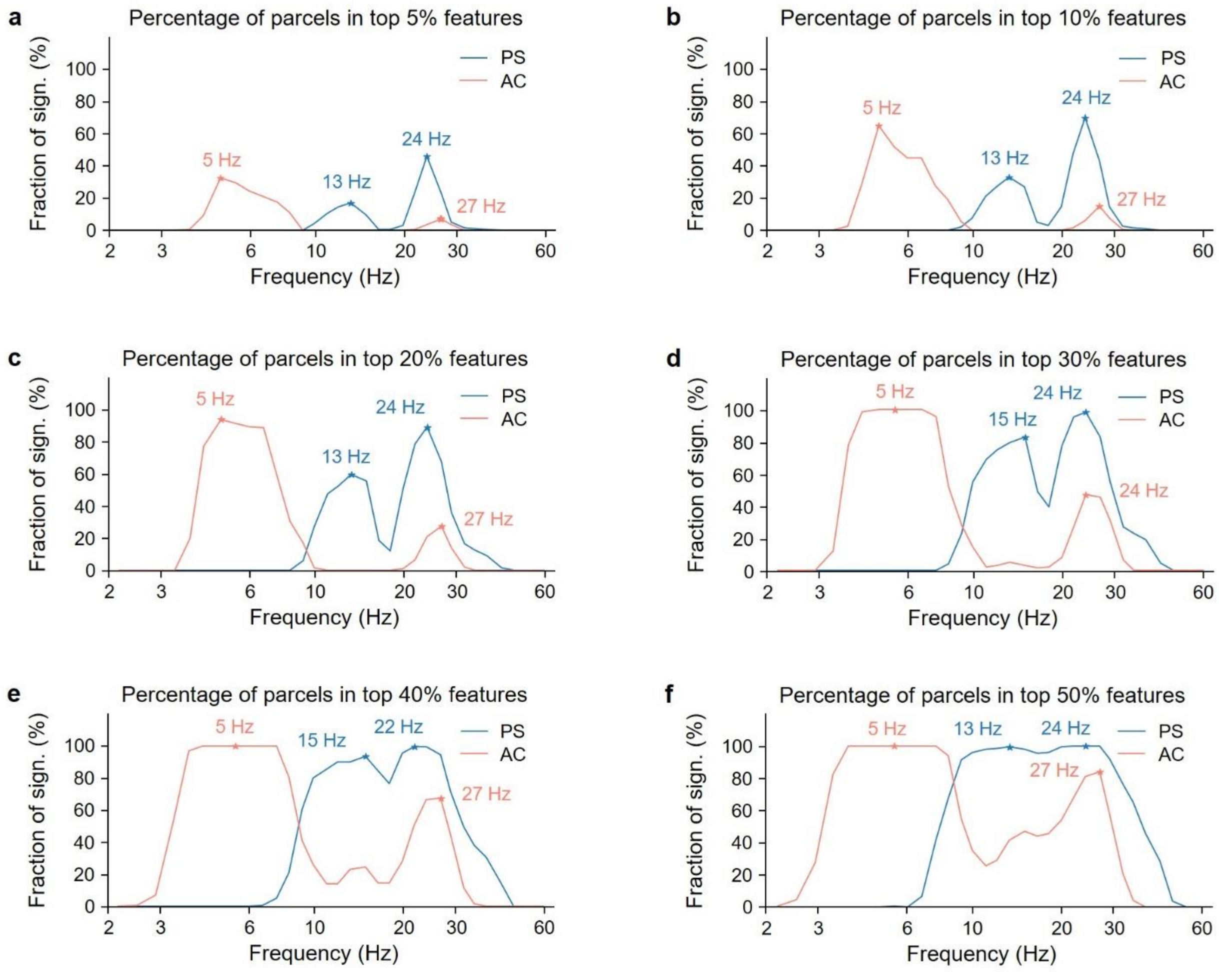

### Supplementary Figure S9

Contributions of amplitude correlation (AC) and phase synchrony (PS) on latent brain-behavior associations. **a** The correlation coefficients between the AC-derived significant latent component and parcel node strength at 5 Hz and 27 Hz. **b** The correlation coefficients between the PS-derived significant latent component and parcel node strength at 13 Hz and 24 Hz. **c** Bar plot of the sum of correlation coefficients from **a** for each subnetwork. **d** Bar plot of the sum of correlation coefficients from **b** for each subnetwork.

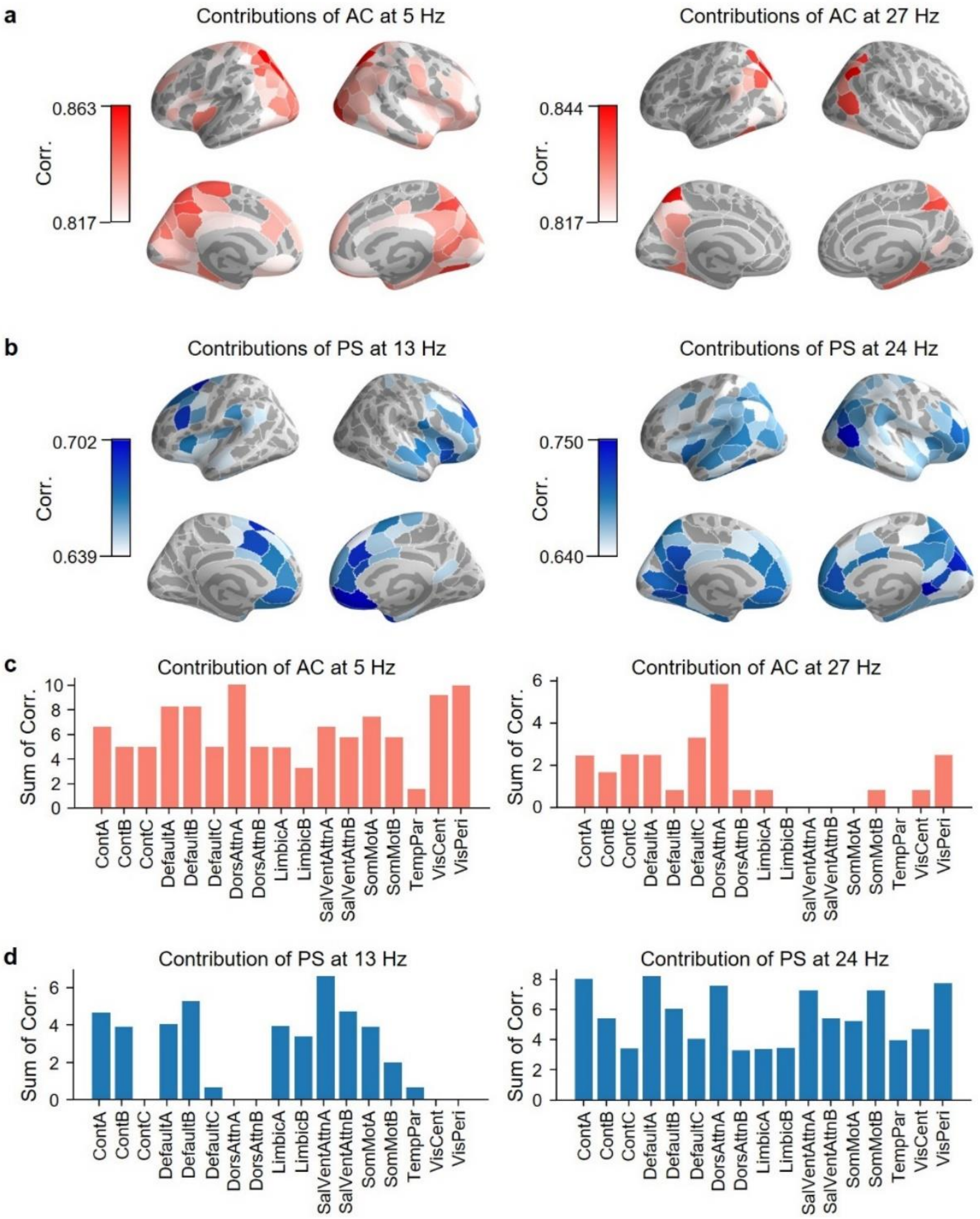

### Supplementary Figure S10

Comparison between males and females on AC and PS for the MDD cohort. The spectrum of graph strength of AC (left) and PS (right) for the MDD cohort (*N*=263). No significant differences in graph strength were found for neither AC and PS between male and female cohorts (Mann-Whitney U test, *p*>0.05, False Discovery Rate (FDR) corrected).

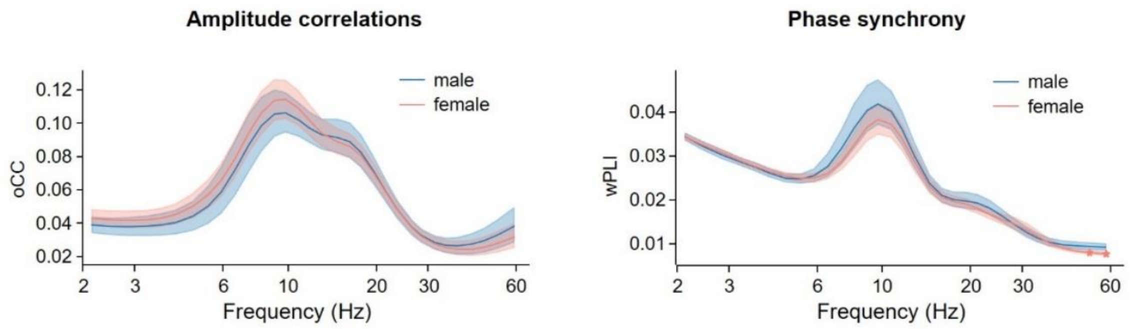

### Supplementary Figure S11

Contributions of symptoms and connectivity node strength on latent brain-behavior associations for the female and male MDD cohorts. For the female cohort (N=163), one latent component was found between PS and symptoms (*p=0.0027*), and for the male cohort (N=87), one latent component was found between AC and symptoms (*p=0.027*). **a** The contribution of symptoms on the two latent components (one from **PS, *p=0.0027*** and one from AC, *p>0.05*) for the female cohort. **b** Fraction of parcels per frequency among the top 10% of features contributing the most to the two latent components for the female cohort. **c** The contribution of symptoms on the two latent components (one from PS, *p>0.05* and one from **AC, *p=0.027***) for the male cohort. **d** Fraction of parcels per frequency among the top 10% of features contributing the most to the two latent components for the male cohort.

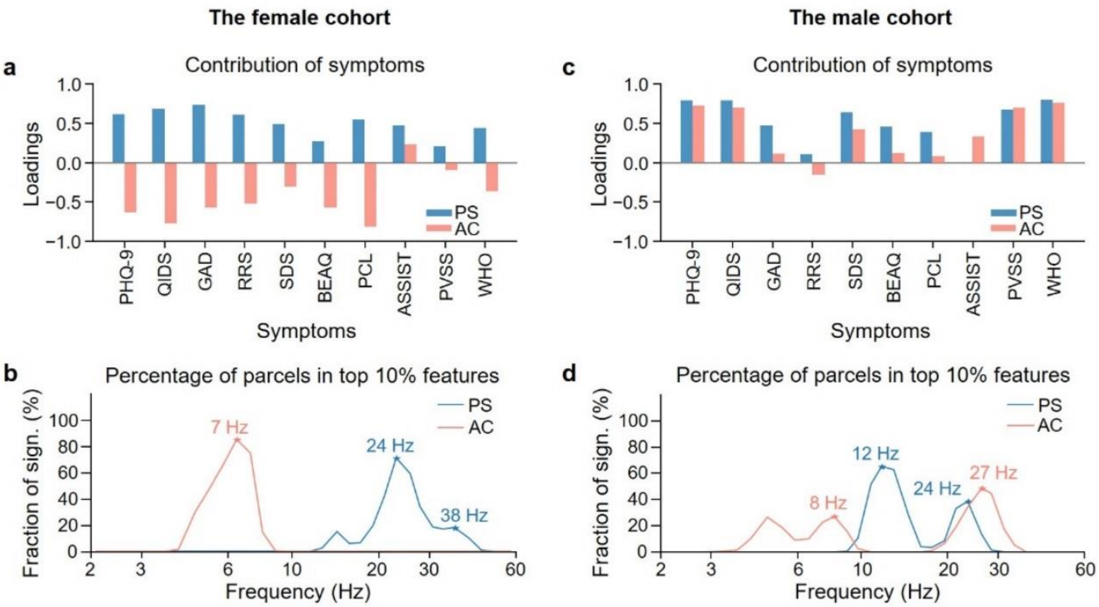

#### Supplementary Figure S12

Age effect on contributions of symptoms and connectivity node strength on latent brain-behavior association. **a** The contribution of symptoms on two latent components (one for AC node strength with *p*=0.0145, and one for PS node strength with *p*=0.0525). **b** Fraction of parcels per frequency among the top 10% of features contributing the most to the two latent components.

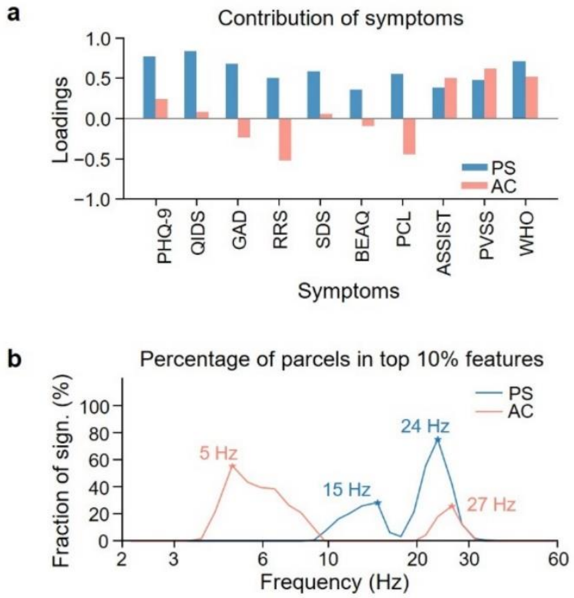

#### Supplementary Figure S13

**a** The number of patients overlapping between clusters obtained with 2 (the first latent component from AC and PS) and 4 (the two latent components from AC and PS) latent components. **b** The Pearson correlations on the similarity matrix obtained with different number of latent components. The patient-patient Euclidean distances were computed with the selected latent components and then transferred to the similarity matrix.

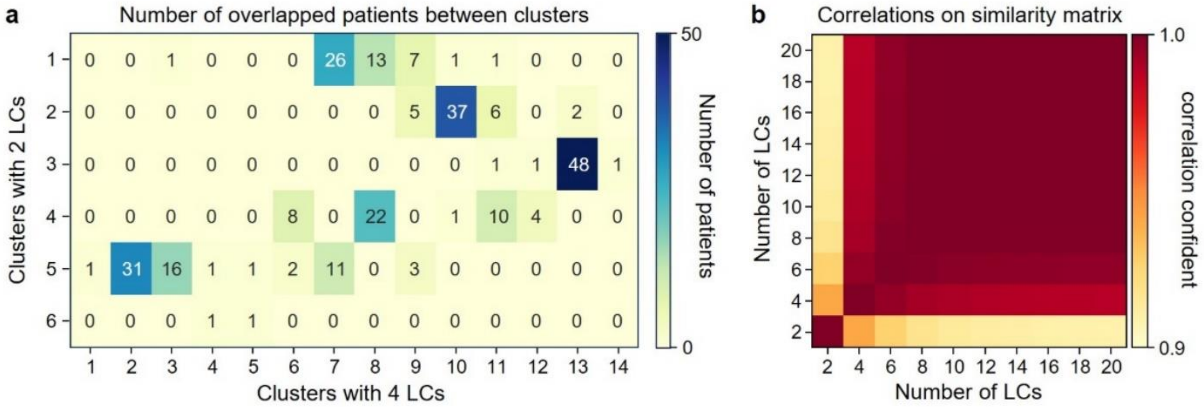

### Supplementary Figure S14

**a** The distribution of node strength latent components for each cluster (left: amplitude correlations, right: phase synchrony). Clusters were compared pairwise with the Mann-Whitney-U test with FDR correction using the Benjamini-Hochberg method. * $p$<0.05, ** $p$<0.01, *** $p$<0.001. **b** Scatter plot of the AC-derived latent component and the PS-derived latent component for each cluster.

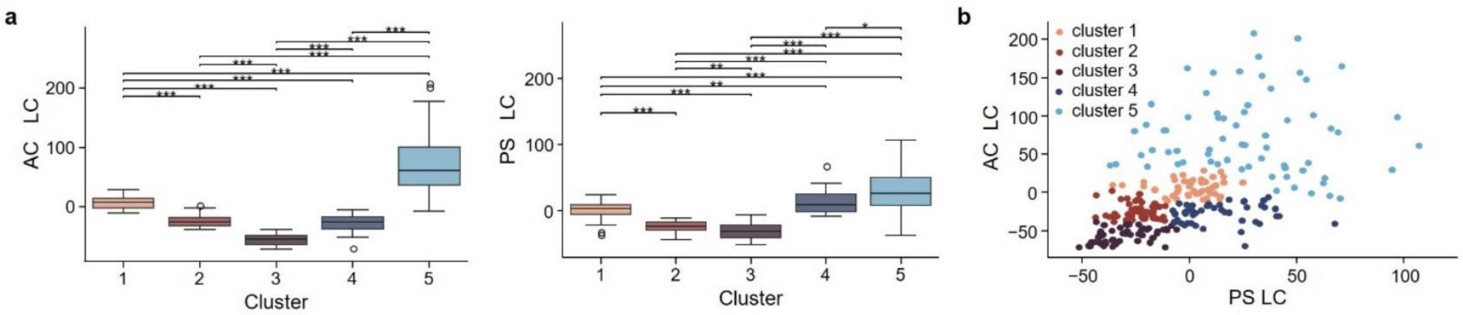

### Supplementary Figure S15

Reproducibility analysis. The entire clustering pipeline was repeated 1,000 times, each time using a randomly selected 80% subset of the full cohort. **a** Each value in the matrix represents the probability that a given pair of patients were assigned to the same cluster across the iterations in which both patients were included. Patients are ordered based on the cluster assignments obtained from the full cohort to facilitate visual interpretation. **b** Probability of reproducibility for each cluster. Bar plots show the mean of reproducibility probability for each cluster, and error bars represent the 2.5th–97.5th percentiles.

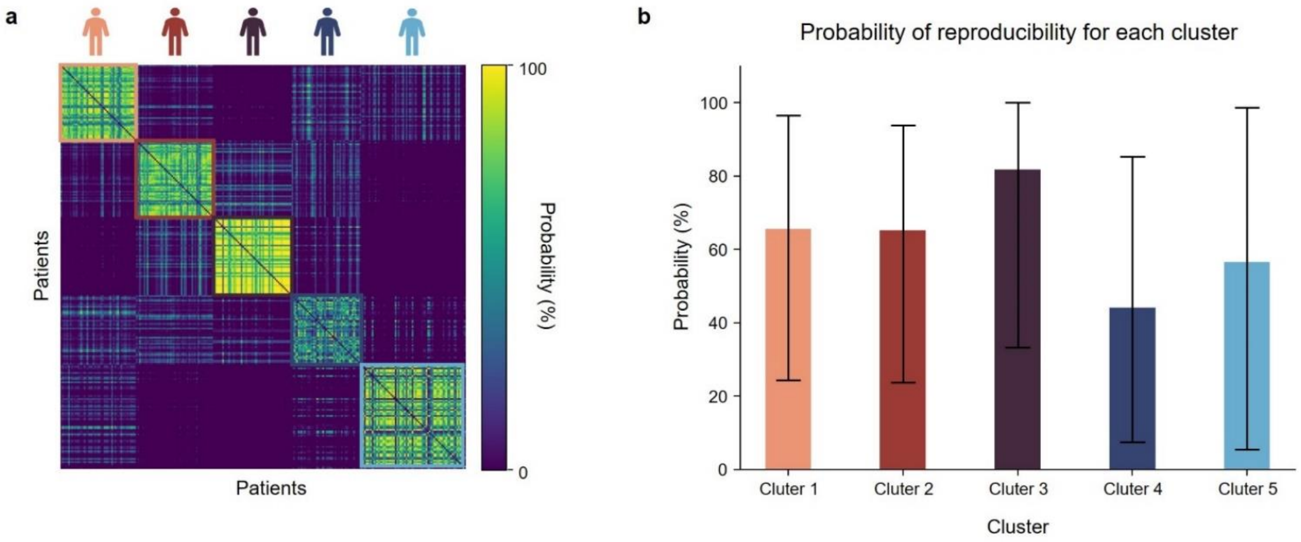

### Supplementary Figure S16

Classification analysis for each cluster. To evaluate the discriminability of each depression phenotype, we trained a series of binary classifiers, one for each cluster, using a one-vs-rest approach. Specifically, for each of the five clusters, we labeled the samples in that cluster as one class and samples from the remaining four clusters as the other class. We then applied a 5-fold cross-validated decision tree classifier to predict cluster membership, yielding classification accuracy for each individual cluster. Bar plot showed the mean classification accuracy for each cluster, and error bars represent the 2.5th–97.5th percentiles across patient in each cluster.

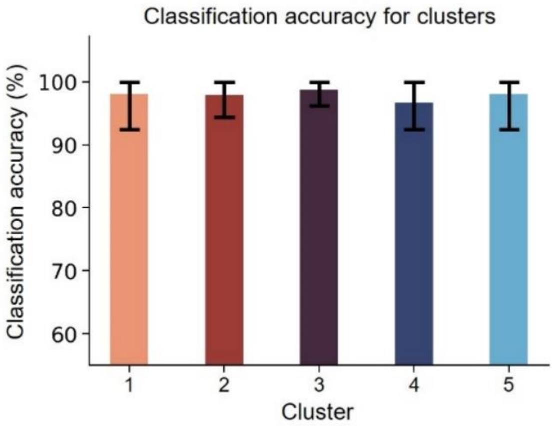

### Supplementary Figure S17

Fraction of significant parcels for AC (**a**) and PS (**b**) node strength for five phenotypes compared with the HC group at *p*<0.05 (Mann-Whitney U test, FDR correction with Benjamini-Hochberg).

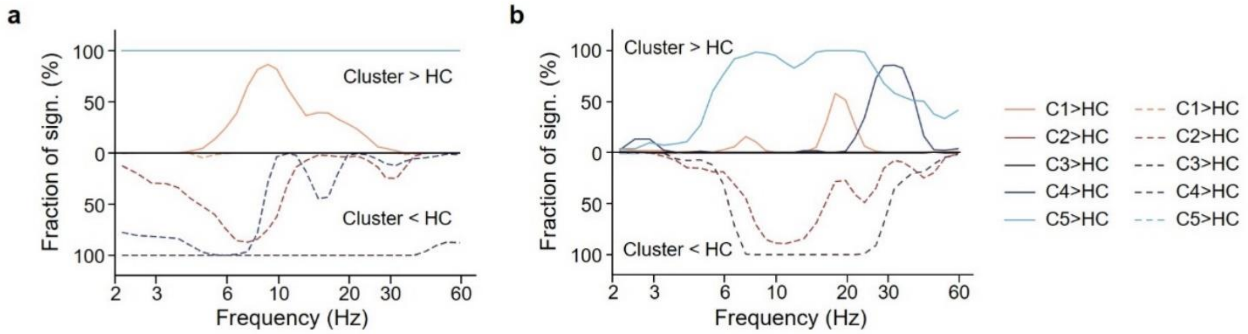

### Supplementary Figure S18

Anatomies of each cluster. **a** Anatomies of averaged AC node strength at 5 Hz across patients in each cluster. **b** Anatomies of averaged AC node strength at 27 Hz across patients in each cluster. **c** Anatomies of averaged PS node strength at 13 Hz across patients in each cluster. **d** Anatomies of averaged PS node strength at 24 Hz across patients in each cluster.

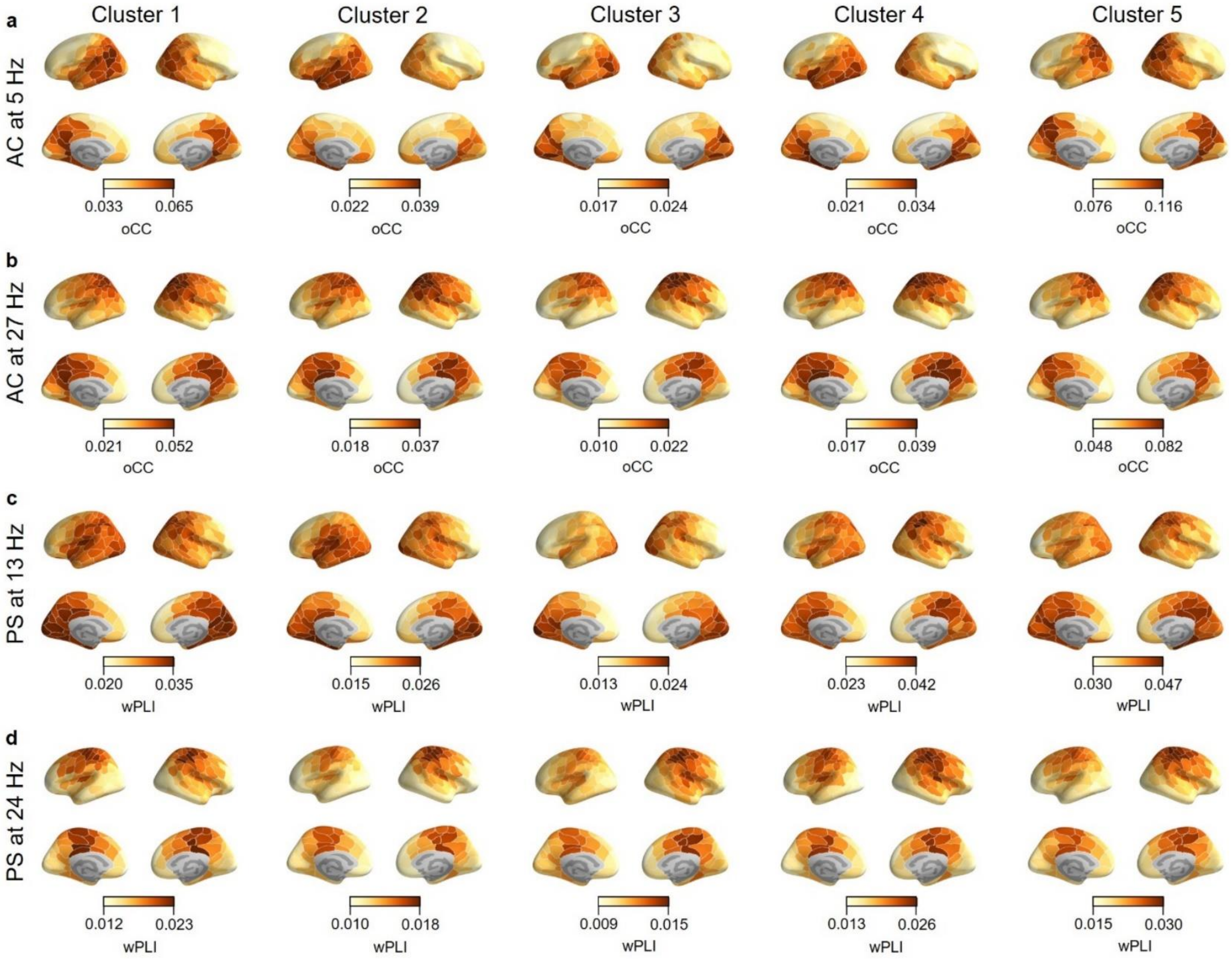

### Supplementary Figure S19

Phenotype variability of AC connectivity strength between parcels at 5 Hz (**a**) and 27 Hz (**b**) and PS connectivity strength between parcels at 13 Hz (**c**) and 24 Hz (**d**). Averaged connectivity strength for each cluster was min-max normalized to the range of 0–1, and then their standard deviations were calculated across clusters.

**a**

Phenotype variability of AC at 5 Hz

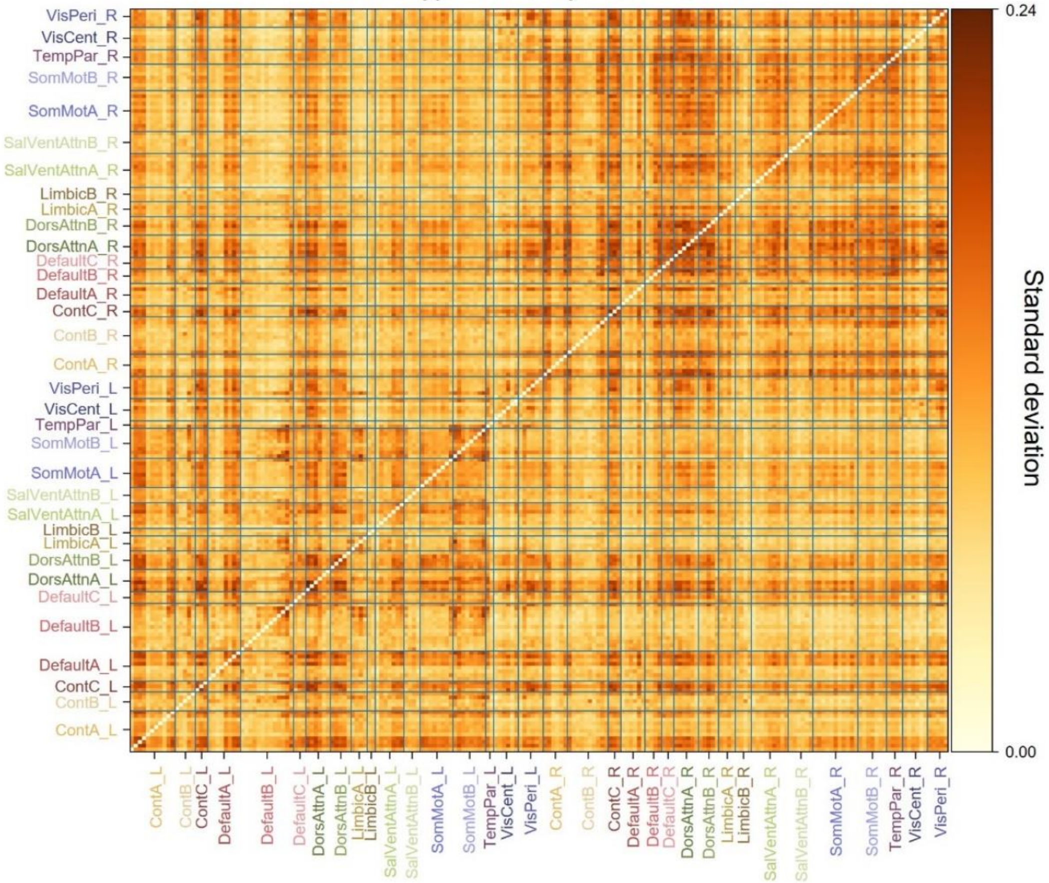


**b**

Phenotype variability of AC at 27 Hz

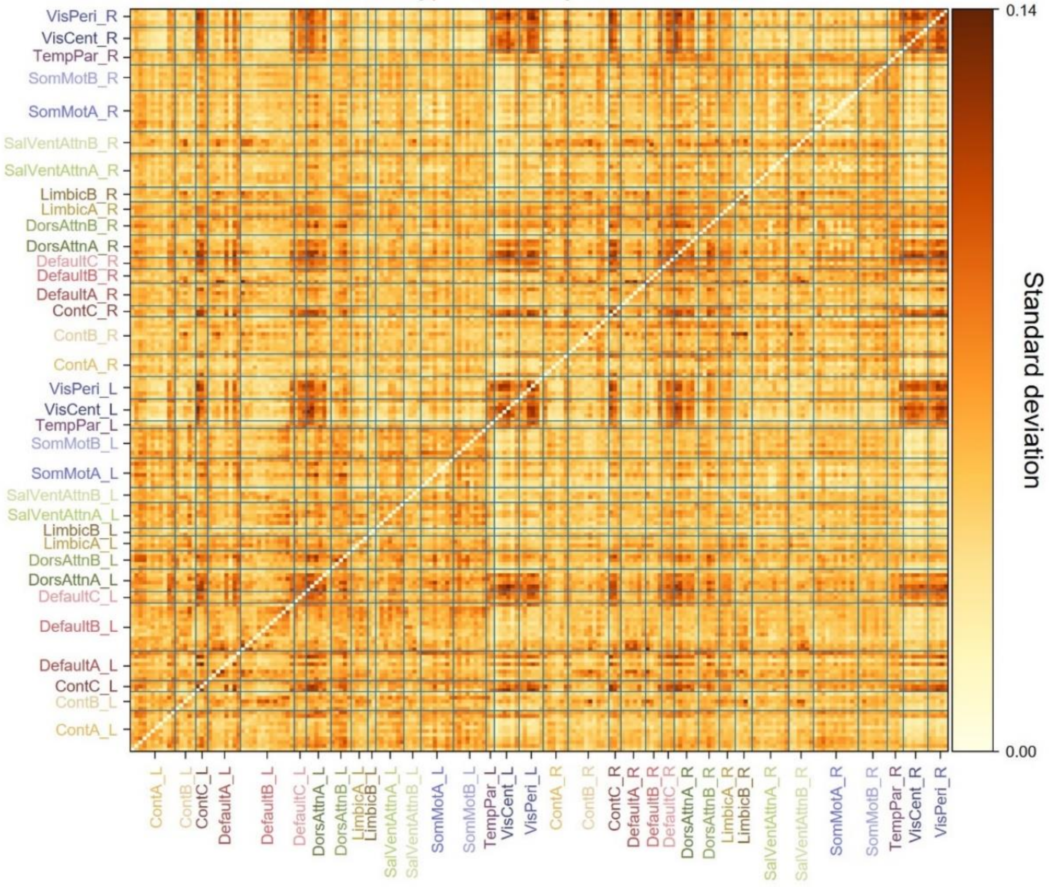

**c**

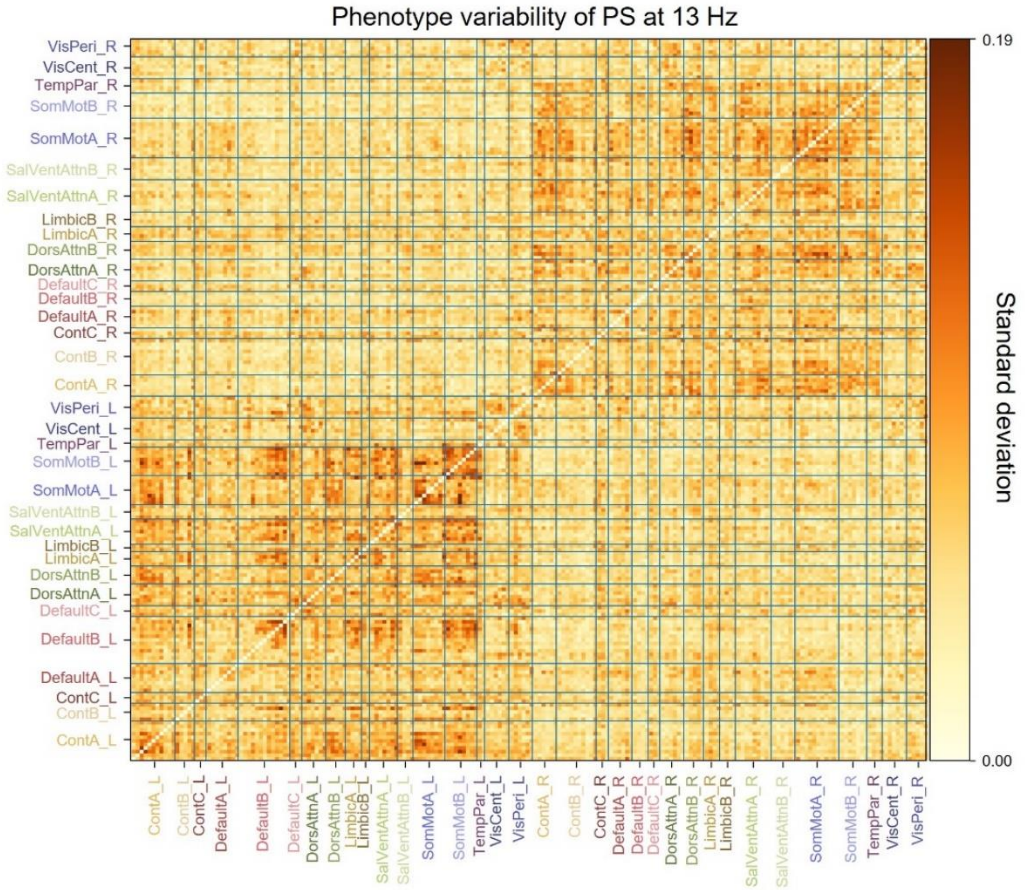
Phenotype variability of PS at 13 Hz
Standard deviation
0.19
0.00
VisPeri_R
VisCent_R
TempPar_R
SomMotB_R
SomMotA_R
SalVentAttnB_R
SalVentAttnA_R
LimbicB_R
LimbicA_R
DorsAttnB_R
DorsAttnA_R
DefaultC_R
DefaultB_R
DefaultA_R
ContC_R
ContB_R
ContA_R
VisPeri_L
VisCent_L
TempPar_L
SomMotB_L
SomMotA_L
SalVentAttnB_L
SalVentAttnA_L
LimbicB_L
LimbicA_L
DorsAttnB_L
DorsAttnA_L
DefaultC_L
DefaultB_L
DefaultA_L
ContC_L
ContB_L
ContA_L

**d**

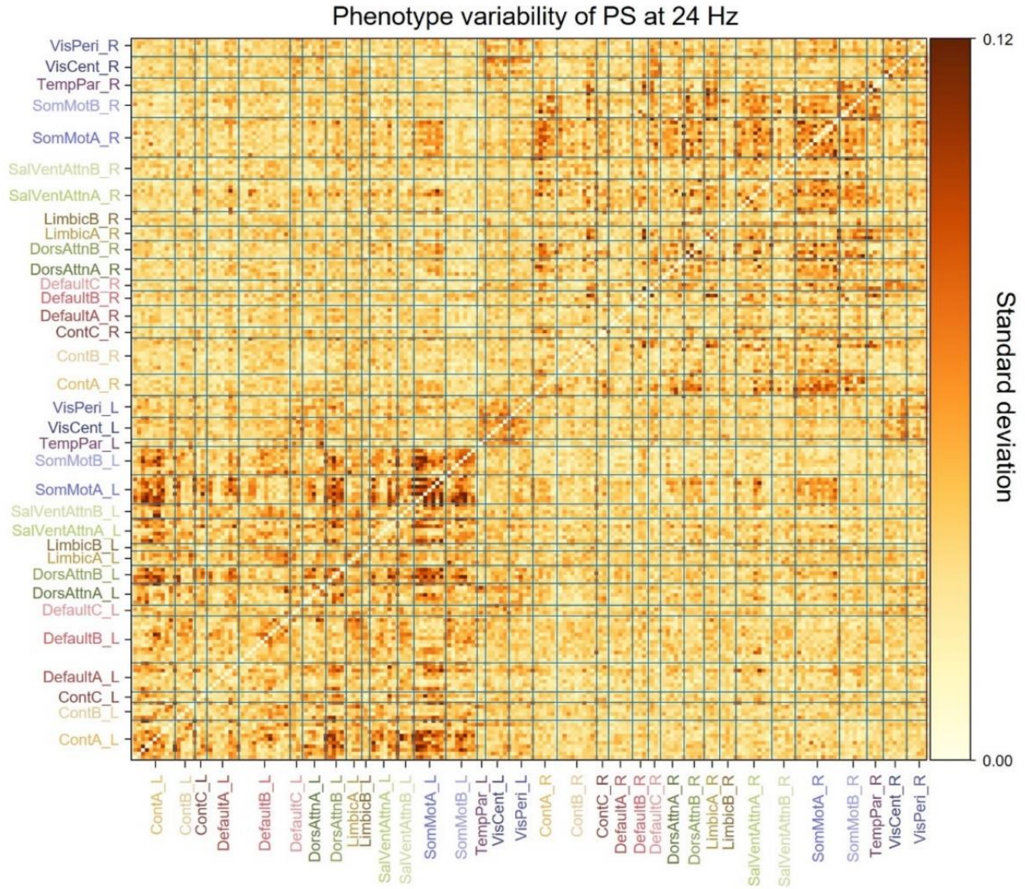
Phenotype variability of PS at 24 Hz
Standard deviation
0.12
0.00
VisPeri_R
VisCent_R
TempPar_R
SomMotB_R
SomMotA_R
SalVentAttnB_R
SalVentAttnA_R
LimbicB_R
LimbicA_R
DorsAttnB_R
DorsAttnA_R
DefaultC_R
DefaultB_R
DefaultA_R
ContC_R
ContB_R
ContA_R
VisPeri_L
VisCent_L
TempPar_L
SomMotB_L
SomMotA_L
SalVentAttnB_L
SalVentAttnA_L
LimbicB_L
LimbicA_L
DorsAttnB_L
DorsAttnA_L
DefaultC_L
DefaultB_L
DefaultA_L
ContC_L
ContB_L
ContA_L

### Supplementary Figure S20

**a** Numbers of males and females in each cluster. No significant differences were found between each pair of clusters (Chi-square test at $p<0.05$, FDR corrected with Benjamini-Hochberg). **b** Age distributions for males and females in each cluster. No significant differences were found between each pair of clusters (the Mann-Whitney U at $p>0.05$, FDR correction with Benjamini-Hochberg).

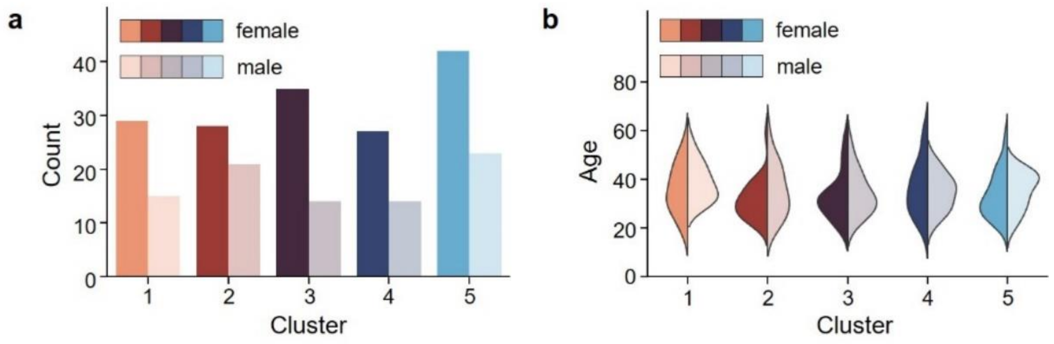
